# Defective HIRA driven chromatin maintenance underlies age related decline in oocyte quality

**DOI:** 10.64898/2026.09.28.755122

**Authors:** Yuki Hatanaka, Yuko Takeda, Max Hannam, Erin E. Cutts, Buhe Nashun, Marian Dore, Ben Moyon, George Young, Pierre-Olivier Estève, Sriharsa Pradhan, Mary Herbert, Petra Hajkova

## Abstract

The rapid fertility decline in women over 35 years of age has been attributed to decreasing oocyte quality. Aging oocytes accumulate DNA damage, chromosome segregation defects and altered epigenetic modifications. Critically, the molecular mechanisms underlying these changes remain poorly understood. Here we uncovered that in the mouse, the HIRA driven H3.3 histone replacement is attenuated during oocyte aging leading to the loss of chromatin homeostasis. The loss of HIRA function is caused by its SUMOylation resulting in the disruption of HIRA-UBN1-CABIN1 complex and its association with chromatin. We further show that preventing HIRA SUMOylation restores H3.3 incorporation and chromatin integrity in aged mouse oocytes leading to improved oocyte maturation and preimplantation development rates. Importantly, similar chromatin homeostasis defect is observed in aging human oocytes pointing towards a conserved process. Our findings provide novel insights into the molecular mechanisms underlying age-related oocyte quality decline and pave the way for clinical interventions.

---

All nuclear processes involving DNA occur in the context of chromatin, which provides not only protection of the genome against genotoxic stress, but critically, through the combination of histone modifications and 3D structure, underpins transcriptional regulation and thus cell fate stability. Recent studies have illuminated the mechanisms underlying the restoration of chromatin structure and the maintenance of epigenetic identity in the context of DNA replication ^1^ ^2^. There is, however, only limited understanding regarding the longitudinal maintenance of chromatin structure and epigenetic information in postmitotic cells, which rely solely on the chromatin assembly pathways operating outside DNA replication^3^. In this context, the development and differentiation of a mammalian oocyte provides a unique system to study chromatin homeostasis. After a final round of DNA replication, developing oocytes enter into meiotic prophase in the mid-gestation foetal gonad (embryonic day (E) 13.5 in the mouse^4^ and gestation week 10 in the human^5^); their development and differentiation then resumes postnatally when individual follicles undergo growth and maturation in response to hormonal stimuli (Fig. 1a). As a consequence, the last round of DNA replication and the final stages of oocyte development can be separated by months (in mouse) or decades (in human).

**Figure 1.**
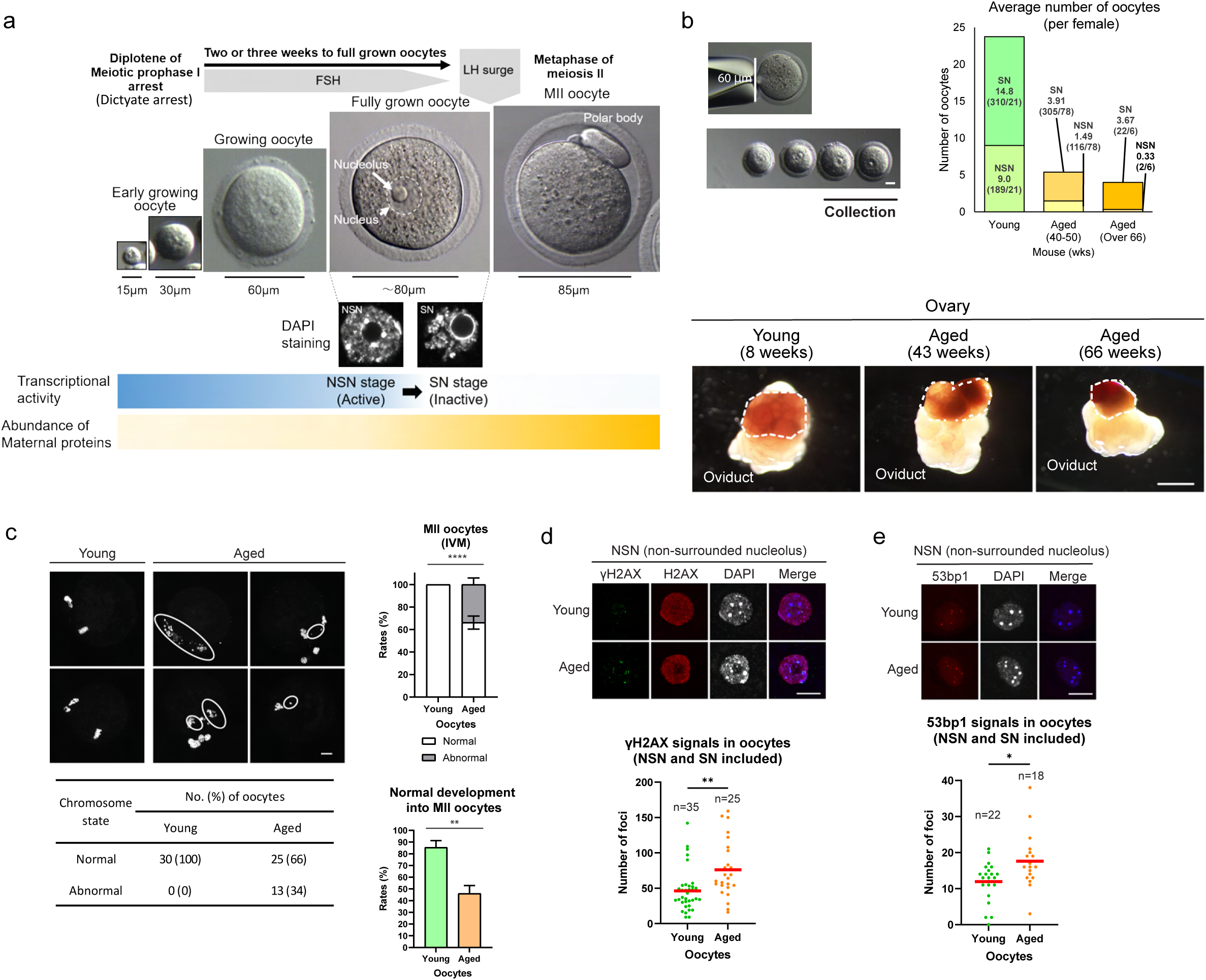
Characterisation of aged oocytes. a. Schematic diagram of oocyte maturation. b. Size selected fully grown oocytes (>60 μm) were used for all analyses (left). Average number of oocytes (NSN and SN stages) per female of a stated age (right). The appearance of young (8 weeks) and aged (43 and 66 weeks) mouse ovaries (bottom). Dotted line delineates the ovary. c. Abnormal chromosome condensation and segregation in aged MII oocytes. Results based on three biological replicates. Bar shows SD. **p < 0.01. ****p < 0.0001. Scale bar, 20μm. “abnormal” refers to the presence of lagging chromosomes. d. γH2AX staining shows significant accumulation of DNA damage in aged oocytes. n=number of analysed oocytes, three biological replicate experiments. **p < 0.01. Scale bar 20μm. Red bars indicate mean. e. IF staining of 53BP1 shows significant accumulation in aged oocytes. n=number of analysed oocytes, *p < 0.01. Scale bar 20μm. Red bars indicate mean.

Oocyte growth in the adult ovary is accompanied by global changes in the transcriptome followed by profound chromatin reorganisation during the final germinal vesicle (GV) stage. When fully grown, oocytes become transcriptionally quiescent and this transition is marked by clustering of heterochromatin around the nucleolus^6^ (non-surrounded nucleolus (NSN) to surrounded nucleolus (SN) oocyte transition, Fig. 1a). This chromatin restructuring underlies an important step in oocyte maturation; compared with the maturation and fertilisation of SN oocytes, the NSN oocytes show lower rates of meiosis resumption and embryo development following fertilisation. This is consistent with differential abundance of transcripts in NSN and SN oocytes^7–9^.

We have previously shown that chromatin in the maturing oocytes is highly dynamic, this even in the absence of active transcription in the SN GV oocytes^10^ (Fig. 1a). We demonstrated that, out of the H3 histone variants, only H3.3 can be incorporated into the chromatin of developing oocytes in the adult ovary and that the maintenance of oocyte chromatin structure is critically dependent on the histone H3.3 chaperone HIRA^10^. Loss of maternal HIRA leads to aberrant transcriptional regulation, appearance of DNA damage, severe oocyte maturation defects and chromosome segregation errors resulting in rapid oocyte loss and female sterility^10,11^.

Given the fact that the arrested oocytes reside in the human ovary for decades before they resume meiosis, it is perhaps not surprising that oocyte quality declines with maternal age. In humans, infertility rates increase rapidly in women over 35 years of age^12^. This correlates with an increase in rate of chromosomal abnormalities (on average ∼ 30% in 35-39-year-olds,∼60% in 40-46-year-olds, but can be even higher depending on the individual)^12–15^. Across many species, including human, this decline in oocyte quality has been linked to numerous factors including progressive decay of sister chromatid cohesion during the dictyate arrest^16–19^ (Fig1a), accumulation of DNA damage, aberrant nuclear and nucleolar architecture, oxidative stress caused by environmental exposure and epigenetic alterations^20–25^. However, the underlying causes, the exact molecular mechanisms and consequently, the routes for potential therapeutic interventions have remained subject of intense debate and investigations.

## Results

### DNA damage and oocyte maturation defects in aged oocytes

To gain a deeper understanding of the oocyte aging process, we focused on the analysis of oocytes isolated from hormonally unstimulated ovaries of 40-50 weeks old C57Bl6/J females. Only fully grown oocytes (>60 µm; see Materials and Methods and Fig. 1b) were included into the analysis to avoid variability due to incomplete maturation. The choice of this aging “window” was driven by several considerations: While the females can still produce healthy offspring, they show clear signs of fertility decline as they are approaching the end of their reproductive lifespan^26,27^ : 1) The ovaries at this age show apparent morphological changes including reduced size and signs of inflammation^28–30^, 2) the numbers of oocytes in the ovaries are severely reduced^26,31^ (Fig. 1b), and 3) when subjected to in vitro maturation (IVM), around 30% of oocytes show chromosome defects (lagging chromosomes or complete failure of chromosome condensation and segregation) upon reaching the MII stage (Fig. 1c). This is similar to the levels observed in human oocytes from women over 35 years of age^12–15^. In addition, and confirming previous findings^32–34^, the oocytes from 40-50 weeks old mouse females show accumulation of γH2AX and 53BP1 foci, a clear sign of persistent DNA damage (Fig 1d,e).

### Aged oocytes show reduced capacity for H3.3 incorporation leading to increased chromatin accessibility

We have previously shown that the chromatin homeostasis in developing oocytes critically relies on the ongoing histone turnover and H3.3 incorporation^10^. Given the longevity of oocytes and the temporal separation of the oocyte development and the last S-phase, we first asked whether this histone replacement can be affected by progressive oocyte age. Injection of mRNAs for H3.1-GFP, H3.2-GFP, and H3.3-GFP (Fig. 2a) confirmed that out of the newly synthesised H3 histones, only H3.3 shows chromatin localisation in GV oocytes, in agreement with our previous study^10^. However, and unexpectedly, the H3.3 incorporation was significantly reduced in the aged oocytes (Fig. 2b). The observed reduced H3.3 incorporation is not due to changes in translation and / or nuclear import in the aged oocytes, as confirmed by microinjection of the mRNA for GFP linked to nuclear localisation signal (NLS-GFP) (Extended Data Fig. 1a,b).

**Figure 2.**
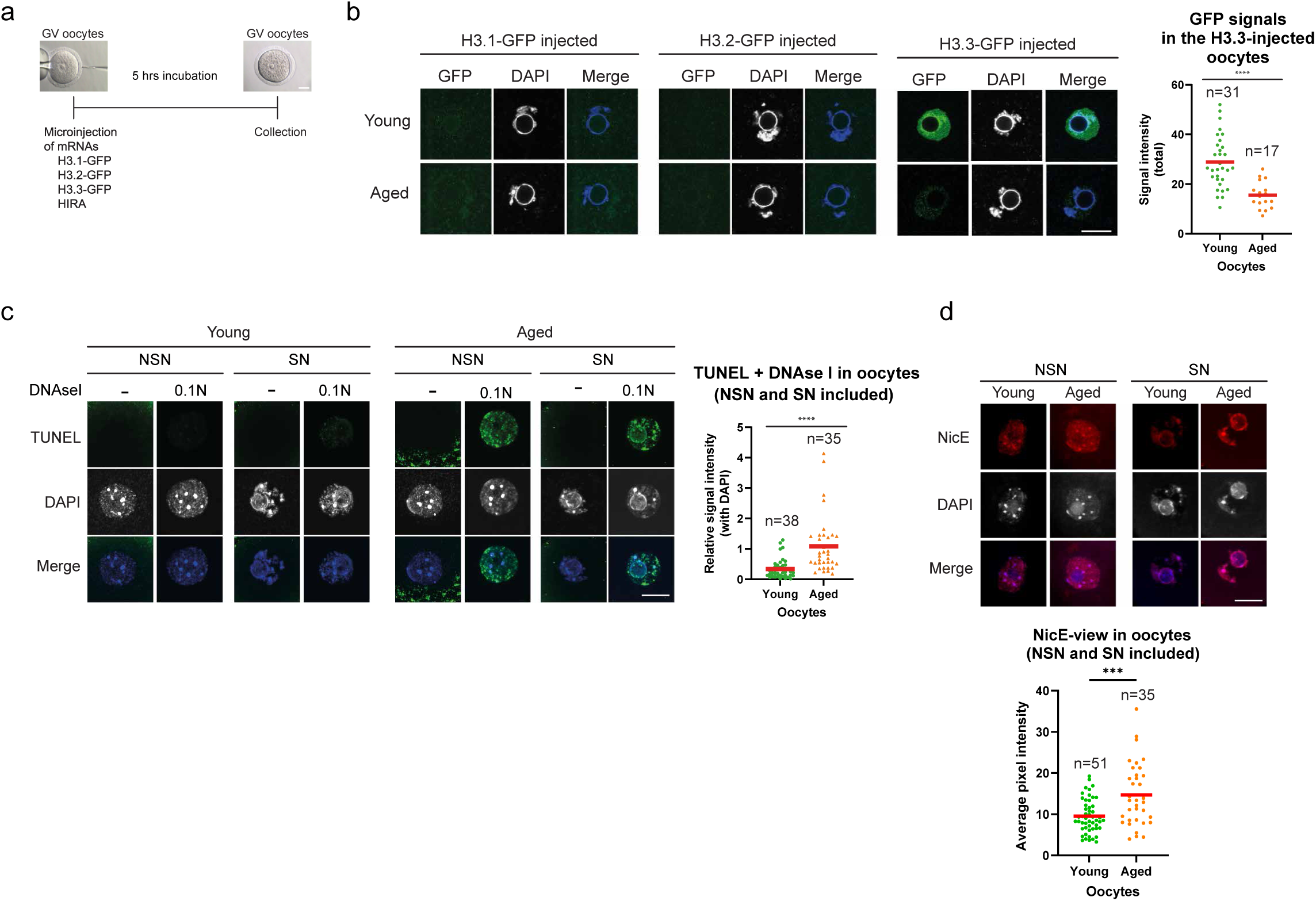
Changes of chromatin dynamics between young and aged oocytes. a. Scheme of mRNA microinjection into GV oocytes. b. Incorporation of newly synthesised H3 variants in young and aged oocytes analysed by immunofluorescence. Quantification of H3.3-GFP incorporation, each data point represents an individual microinjected oocyte. n=number of analysed oocytes, ****p < 0.0001. Red bars indicate mean. Scale bar 20μm. c. DNAseI treatment followed by TUNEL assay was carried out on young and aged NSN and SN GV oocytes to compare global chromatin accessibility. n=number of analysed oocytes (NSN and SN), ****p < 0.0001. Red bars indicate mean. Scale bar 20μm. d. NicE-view documents increased DNA accessibility in aged oocytes (NSN and SN stages). n=number of analysed oocytes, ***p < 0.001. Red bars indicate mean. Scale bar 20μm.

Oocyte specific deletion of the H3.3 chaperone HIRA and the consequent lack of H3.3 incorporation leads to nucleosomal loss and increased DNA accessibility^10^. We hence used DNAseI treatment followed by a fluorescent TUNEL assay^10^ to examine whether, in line with the reduced H3.3 incorporation, chromatin accessibility differs in individual young and aged oocytes. Fluorescent TUNEL signal was readily detectable in the aged oocytes (both NSN and SN stages) even at low DNAseI concentrations, in contrast to only a background signal observed in the young oocytes (Fig. 2c, Extended Data Fig. 1c,d). These findings were additionally corroborated using an orthogonal approach “NicE-view” that allows for visualisation of regions with accessible chromatin^35^. Also this approach confirmed increased chromatin accessibility in the aged (both NSN and SN) oocytes (Fig. 2d and Extended Data Fig. 1e,f).

### SUMO2/3 modification of HIRA in aged oocytes leads to decreased interaction with Cabin1 and reduced chromatin association

We next considered whether the reduced H3.3 incorporation might be caused by transcriptional downregulation of the HIRA histone chaperone or its interacting partners^39^ using single oocyte RNASeq analysis of young and aged (40-50 weeks) GV oocytes. Notably, the analysis of the included ERCC RNA spikes revealed higher amplification in the aged oocyte libraries documenting lower RNA content in the aged oocytes (Extended Data Fig. 2a). We further probed this using EU incorporation that confirmed lower overall transcription in the aged oocytes (Extended Data Fig. 2b). NSN oocytes show high transcriptional activity followed by transcriptional quiescence at the SN stage^36^. As expected, lower EU incorporation was observed in the aged NSN oocytes, while no EU incorporation was detected in either young or old SN oocytes (Extended Data Fig. 2b).

Our single oocyte RNASeq revealed only very limited transcriptional differences between young and aged oocytes identifying 153 differentially expressed genes (73 upregulated, 80 downregulated genes, young vs aged, padj<0.05) (Extended Data Fig.2c-f and Extended Data Table 1) and only a few changes in the expression of transposable elements (TEs, Extended Data Fig. 2g,h and Extended Data Table 2). The reduced number of differentially expressed genes in comparison with previous studies^27,37^ is likely due to an earlier age point (11 vs 14 months of age), size selection of oocytes and importantly, lack of hormonal stimulation that has been previously shown to contribute to oocyte abnormalities^38^.

Relevant to our earlier findings, we noted that both Cabin1 and Ubn1 – interaction partners in the HIRA chaperone complex^39^ – were significantly, albeit moderately downregulated in the aged oocytes (Extended Data Fig. 2f,i). These differences were however not confirmed at the protein level, as Cabin1, UBN1, as well as ASF1a showed no apparent differences in abundance between young and aged (40-50 weeks) oocytes (Extended Data Fig. 3a).

To the contrary (and in discordance with our RNASeq analysis, Extended Data Fig. 2i), the signal for HIRA protein was significantly reduced in the aged oocytes (Fig.3a and Extended Data Fig. 3c for antibody specificity). Surprisingly, this was consistently observed only when using an anti-HIRA monoclonal antibody, while equal signal intensity could be detected by the polyclonal anti-HIRA antibody raised against the full-length HIRA protein (Fig. 3b and Extended Data Fig. 3c for antibody specificity). These results suggested the possibility that the epitope recognised by the HIRA monoclonal antibody (Extended Data Fig. 6a) might be masked by a posttranslational modification specifically in the aged oocytes and this could be linked to the observed altered function of the HIRA complex.

**Figure 3.**
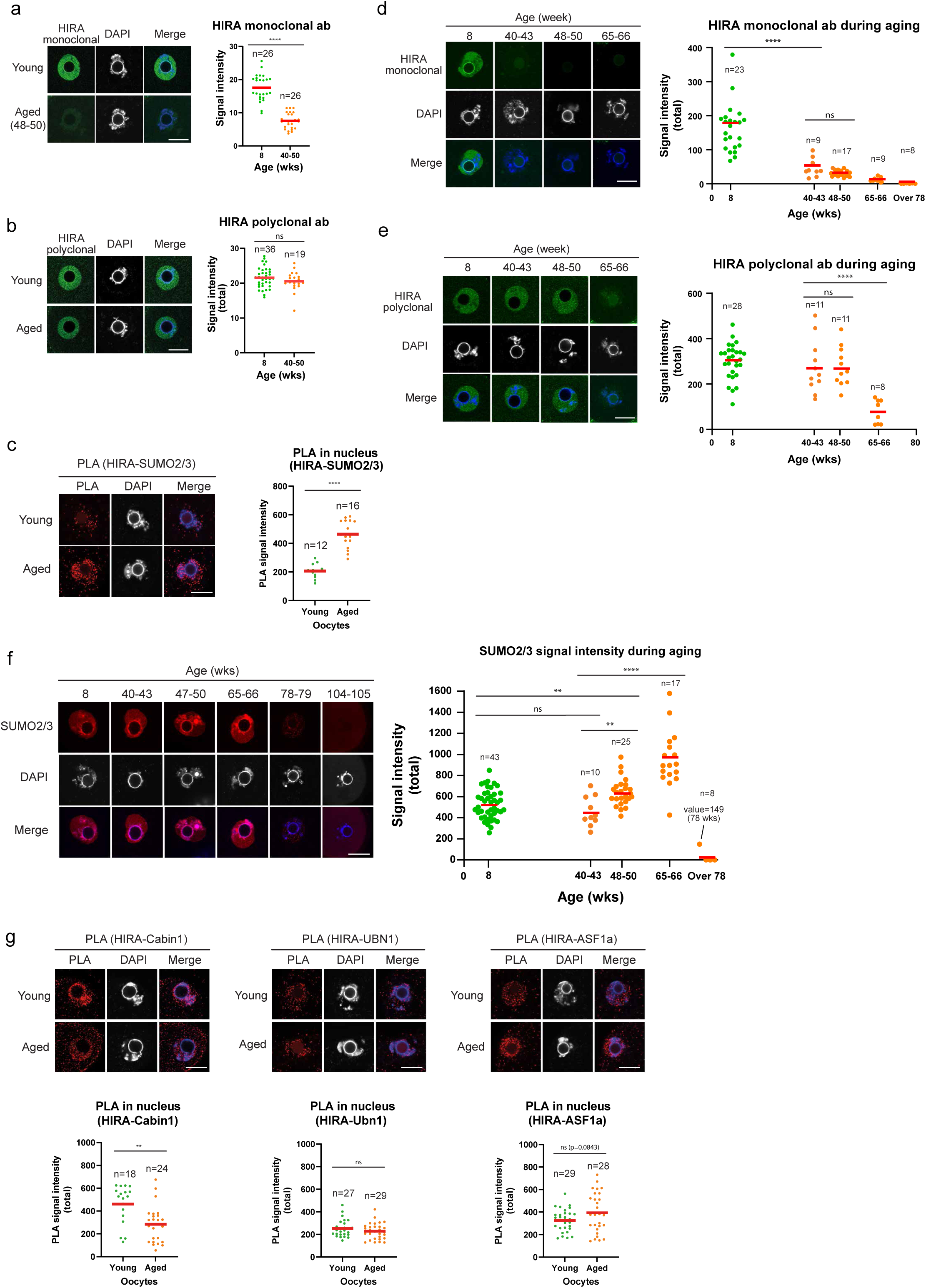
SUMOylation of HIRA and its effect on HIRA complex. a. Immunofluorescence staining using monoclonal HIRA antibody shows reduced signal in aged oocytes. n=number of analysed oocytes, ****p < 0.0001. Red bars indicate mean. Scale bar 20μm. b. Immunofluorescence staining using polyclonal HIRA antibody, note that similar signal is detected between young and aged oocytes. n=number of analysed oocytes, ns – not significant. Red bars indicate mean. Scale bar 20μm. c. PLA between HIRA and SUMO2/3 in SN oocytes. n=number of analysed oocytes, ****p < 0.0001. Red bars indicate mean. Scale bar 20μm. d. Immunofluorescence signal for HIRA monoclonal antibody across oocyte aging (>78 weeks – contains samples from 78-104 weeks old females). n=number of analysed oocytes, ****p < 0.0001, ns – not significant. Red bars indicate mean. Scale bar 20μm. e. Immunofluorescence signal for HIRA polyclonal antibody across oocyte aging. The signal decrease was observed at an advanced age of 65-66 weeks. n=number of analysed oocytes, ****p < 0.0001, ns – not significant. Red bars indicate mean. Scale bar 20μm. f. Immunofluorescence signal for SUMO2/3 reveals progressive accumulation followed by a dramatic loss after week 78 (>78 weeks – contains samples from 78-104 weeks old females). n=number of analysed oocytes, **p < 0.01, ****p < 0.0001, ns – not significant. Red bars indicate mean. Scale bar 20μm. g. PLA between HIRA and other components of the HIRA complex performed in SN oocytes. The interaction between HIRA and Cabin1 is significantly reduced in aged oocytes. n=number of analysed oocytes, **p < 0.01, ns – not significant. Red bars indicate mean. Scale bar 20μm.

Closer investigation revealed that the part of the HIRA protein used for the production of monoclonal antibody harbours several predicted SUMOylation sites (GPS-SUMO 2.0 database - see Materials and Methods; and Extended Data Fig. 6a,b) with HIRA previously reported to be associated with SUMO^40–42^ .

Oocytes have been previously shown to express SUMO1 and SUMO2/3. We confirmed that the signal for SUMO1 is mainly localised at the nuclear membrane (Extended Data Fig. 3d, Extended Data Fig. 4g for antibody specificity), in agreement with a previous report^43^. To the contrary, abundant signal for SUMO2/3 was readily detectable in the nucleus of both young and aged GV oocytes prompting us to focus on the possible relationship between HIRA and SUMO2/3.

As oocytes are not easily amenable to biochemical analyses, we set out to probe whether HIRA can be directly modified by SUMO2/3 *in vitro* and in cultured mouse pluripotent embryonic stem cells (mESCs). We overexpressed Flag-HIRA in wild-type (wt) mESCs (Extended Data Fig. 4a,b), or in mESCs stable expressing HA-SUMO2Q87R (Extended Data Fig. 4e) and were able to detect the presence of SUMO2/3 on HIRA following immunoprecipitation (IP) (Extended Data Fig. 4b,c,e,f, and Extended Data Fig. 4g for SUMO detection specificity). In addition, we also confirmed that HIRA can be modified by SUMO2 using an *in vitro* SUMOylation assay (Extended Data Fig. 4d,g).

Having established that HIRA can be modified by SUMO2/3, we tested the possible interaction in the oocytes using the Proximity ligation assay (PLA) (Extended Data Fig. 3e). As a control, we first assessed the PLA assay functionality and specificity in oocytes using microinjection of mRNAs for either H3.1-GFP or H3.3-GFP followed by PLA with anti-GFP and polyclonal anti-HIRA antibodies (Extended Data Fig. 3f). This confirmed specific interaction between HIRA and H3.3 and lack of interaction between HIRA and H3.1, thus validating this assay in the oocytes. Subsequent PLA using anti-HIRA (polyclonal) and anti- SUMO2/3 antibodies revealed significant accumulation of the signal specifically in the aged oocytes (Fig. 3c), supporting our hypothesis of aged related HIRA SUMOylation.

As the decline in female fertility is a protracted process, we wondered about the kinetics of the observed changes. Interestingly, the initial loss of monoclonal HIRA antibody signal, clearly detectable from 40 weeks of age (Fig. 3d), is followed by the overall loss of the HIRA protein detectable by polyclonal antibody (from about week 65, Fig. 3e) – the timing of the HIRA protein loss coincides with the reported loss of fertility in C57Bl6 females (majority of females are infertile after 12 months^26^). We also observed a progressive accumulation of signal for SUMO2/3 that peaks at about the same time (65-66 weeks) before disappearing in the few oocytes surviving in the ovaries of females older than over 78 weeks (78-104 weeks, Fig. 3f).

It is known that SUMO2/3 modification of target proteins can induce conformation changes leading to disruption of interaction with binding partners^44,45^. In agreement with this, we found that the interaction of SUMO2/3 with HIRA coincides with a decreased interaction between HIRA and Cabin1, in the absence of significant differences in the interactions between HIRA and other partners in the complex: Ubn1 and Asf1a (Fig. 3g).

In addition, and in agreement with altered functionality of the HIRA complex, we found reduced association of Cabin1 and Ubn1 with chromatin specifically in the aged oocytes (Extended Data Fig. 3b).

To further investigate the potential impact of HIRA SUMOylation on the HIRA complex, we generated structural predictions using AlphaFold (see Materials and Methods and Extended Data Fig. 5). We obtained a confident AlphaFold3 model of the HIRA complex and interfaces, and mapped the putative SUMOylation sites of HIRA (Extended Data Fig. 6a,b). No SUMOylation sites were found in the HIRA interfaces with ASF1a, UBN1, H3.3 and H4, consistent with the experimental results indicating that ASF1a and UBN1 are still able to bind when HIRA is SUMOylated (Fig. 3g). However, the AlphaFold model predicted that SUMOylation could interfere with the HIRA/Cabin1 interaction in two ways: 1) residue HIRA K1015 lies near the HIRA/Cabin1 interface, hence SUMOylation is likely to disrupt this interaction. 2) residue K807 lies at the trimerization interface of HIRA; SUMOylation of this site could would affect the trimerization, which is necessary for Cabin1 binding^46^ (Extended Data Fig. 5).

### Mutated HIRA restores H3.3 incorporation and chromatin defects in the aged oocytes

To probe the functional relationship between the SUMOylation of HIRA and the observed chromatin defects in the aged oocytes, we generated a HIRA construct with amino acid substitutions at predicted SUMOylation sites (HIRAmut, Extended Data Fig. 6a,b). We also verified that the expression of the mutated HIRA recapitulates localisation of the endogenous protein (Extended Data Fig. 6c) and the level of expression is comparable in young and aged oocytes (Extended Data Fig. 6d). Microinjection of the HIRAwt into aged oocytes recapitulated increased SUMO2/3 association (Fig. 4a,b), reduced interaction between HIRAwt and Cabin1 (Fig. 4c), loss of reactivity with anti-HIRA monoclonal antibody (Fig. 4d) and reduced association of HIRAwt and of Cabin1 with chromatin (Fig. 4d); suggesting that the injected HIRAwt is subject to the same posttranslational regulation as the endogenous protein. In contrast, the association with SUMO2/3 in aged oocytes was abolished when mutated HIRA (HIRAmut) was microinjected (Fig. 4b). This also improved HIRA association with Cabin1 (Fig. 4c), reactivity with anti-HIRA monoclonal antibody and HIRA chromatin association (Fig. 4d).

**Figure 4.**
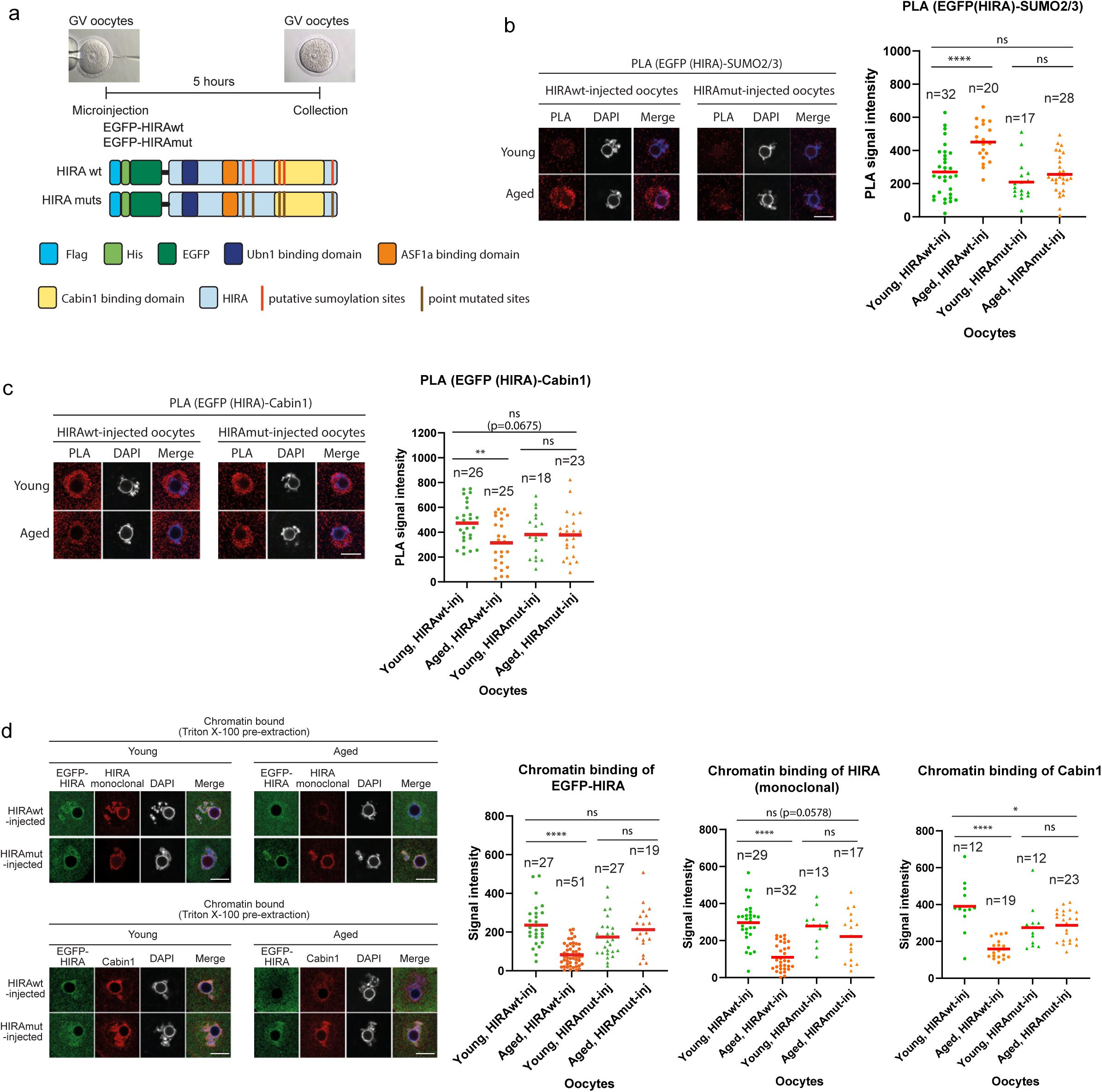
Mutation of SUMO sites restores HIRA and Cabin1 association with chromatin in aged oocytes. a. Microinjection of HIRA (wt and mutated) into GV oocytes. Red lines depict the predicted HIRA SUMOylation sites. EGFP was used for EGFP-HIRA detection in IF and PLA analyses. Injected oocytes were collected 5 hours after microinjection to perform further experiments. b. PLA between HIRA and SUMO2/3 in the HIRAwt- (left) and HIRAmut- (right) injected oocytes. n=number of analysed oocytes, ****p < 0.0001, ns – not significant. Red bars indicate mean. Scale bar 20μm. c. PLA between HIRA and Cabin1 in the HIRAwt- (left) and HIRAmut- (right) injected oocytes. n=number of analysed oocytes, **p < 0.01, ns – not significant. Red bars indicate mean. Scale bar 20μm. d. Chromatin binding of EGFP-HIRA following Triton X-100 pre-extraction prior to fixation. n=number of analysed oocytes, *p < 0.05, ****p < 0.0001, ns – not significant. Red bars indicate mean. Scale bar 20μm.

We next asked whether the expression of mutated HIRA (HIRAmut) could restore the H3.3 incorporation defect observed in the aged oocytes. Although microinjection of the HIRAwt mRNA was not able to rescue the H3.3 incorporation defect, H3.3 incorporation in the aged oocytes was restored upon microinjection of HIRAmut mRNA (Fig. 5a,b). This also restored normal chromatin accessibility (decreased DNAseI sensitivity) and reduced the number of γH2AX foci (DNA damage) to the level observed in the young oocytes (Fig. 5c,d, Extended Data Fig. 7a,b), unequivocally confirming a direct mechanistic link between SUMO modification of HIRA, H3.3 incorporation defect and the observed aberrant chromatin structure.

**Figure 5.**
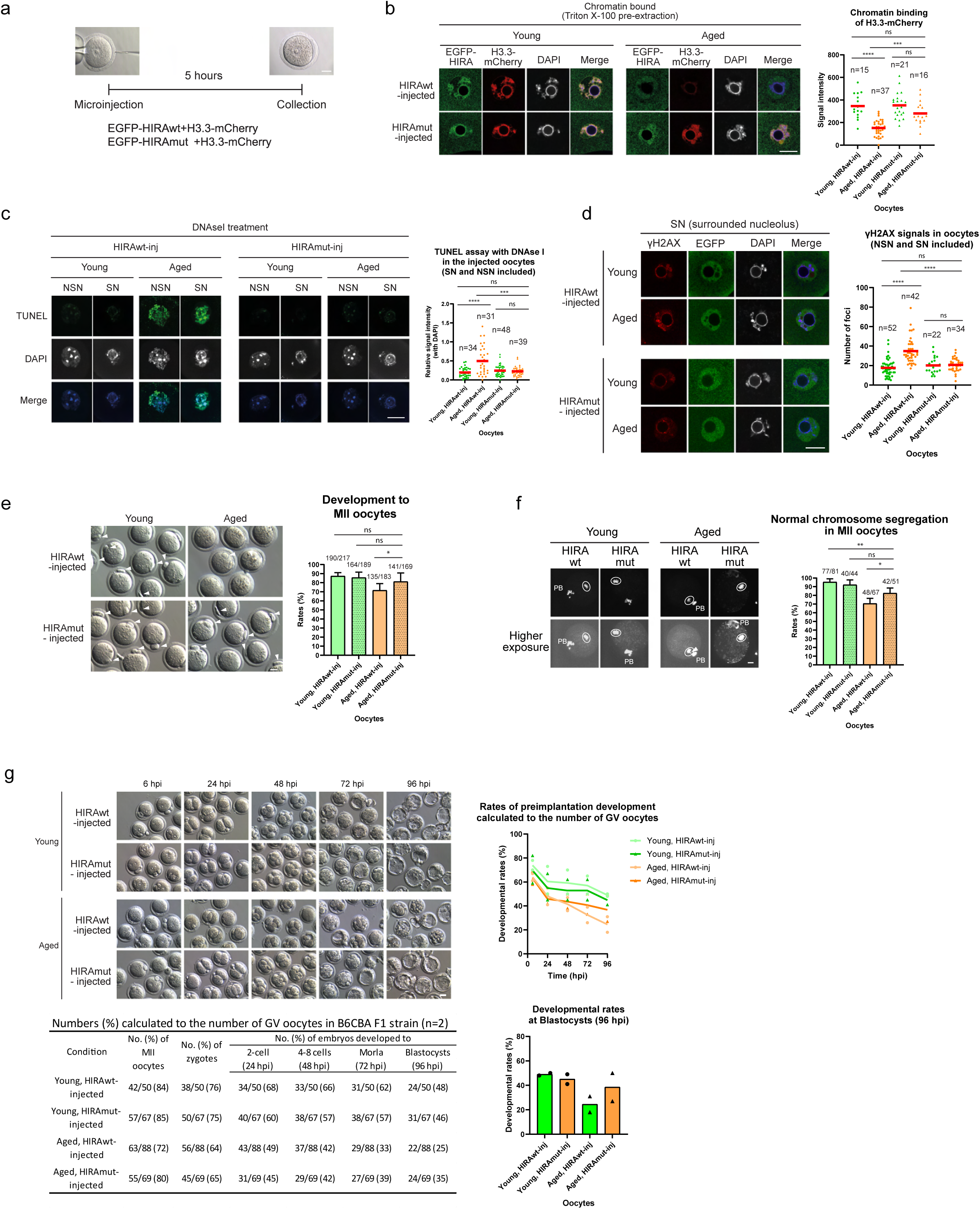
Microinjection of mutated HIRA provides functional rescue in aged oocytes. a. Microinjection scheme to investigate the effect of mutated HIRA on the chromatin deposition of newly synthesised H3.3. b. Chromatin binding of EGFP (HIRA) and mCherry (newly synthesised H3.3) in the HIRAwt (top) and the HIRAmut (bottom) injected oocytes. n=number of analysed oocytes, ***p < 0.001, ****p < 0.0001, ns – not significant. Red bars indicate mean. Scale bar 20μm. c. Z-Stacked images of TUNEL assay following DNAseI treatment in HIRAwt- (right) and HIRAmut-(left) injected oocytes. n=number of analysed oocytes, ***p < 0.001. ****p < 0.0001, ns – not significant. Red bars indicate mean. Scale bar 20μm. d. IF staining of γH2AX in HIRAwt- (right) and HIRAmut-(left) injected oocytes shows significant improvement in aged oocytes following injection of HIRAmut . n=number of analysed oocytes, ****p < 0.0001, ns – not significant. Red bars indicate mean. Scale bar 20μm. e. Efficiency of oocyte maturation to MII stage. In vitro maturation (IVM) of the HIRAwt- or HIRAmut- injected oocytes. Numbers above each bar indicate the number of MII oocytes over the total number of injected oocytes. Arrowheads depict 1^st^ polar body. Scale bar 30μm. *p < 0.01, ns – not significant. Bar shows SD. f. Chromosome condensation and segregation in MII oocytes. Numbers above each bar indicate the number of MII oocytes over the total number of injected oocytes. Arrowhead shows the segregation error. Circles show mitotic chromosomes. PB, polar body. Scale bar 20μm. **p < 0.01. *p < 0.05. ns - not significant. Bar shows SD. g. Preimplantation development following IVM-IVF of HIRAwt or HIRAmut injected young and aged oocytes. Efficiency calculated based on a number of oocytes isolated from the ovary. Scale bar 30μm. Two biological replicates were performed.

Aged oocytes show reduced maturation to MII oocytes due to incomplete chromosome condensation and segregation defects (Fig. 1c). Microinjection of HIRAmut mRNA resulted in improved maturation rate of the aged oocytes relative to the control (83% in HIRAmut vs 74% in HIRAwt), as judged by the emergence of the first polar body (Fig. 5e, Extended Data Fig. 7c). We also observed a significantly increased rate of normal chromosome condensation and segregation in aged oocytes injected with HIRAmut (82% in HIRAmut vs 72% in HIRAwt) (Fig. 5f, Extended Data Fig. 7d), leading to an overall (combined) improvement of the rate of aged oocyte development (Extended Data Fig. 7e). To further assess the quality of the HIRA mut injected oocytes we decided to test their developmental potential. As C57Bl/6J embryos suffer from 2cell stage developmental block *in vitro*^51^, we switched to B6CBA F1 mouse strain. The developmental rate was calculated by dividing the number of oocytes/embryos by the total number of oocytes collected from ovaries. When subjected to IVM followed by IVF, HIRAmut injected B6CBA F1 aged oocytes showed better rates of preimplantation development (35% HIRAmut vs 25% HIRAwt, Fig. 5g, Extended Data Table 3), indicating that improved maturation and quality of aged oocytes could be achieved by improving chromatin structure through microinjection of HIRA that cannot undergo SUMO modification.

### Inhibition of SUMOylation improves oocyte maturation and early embryo development by restoring normal HIRA function in aged oocytes

Our results prompted us to investigate whether we could improve the quality of the aged oocytes using recently identified potent inhibitors of SUMOylation^47–49^ . We tested two specific inhibitors: 1) 2-d08, an inhibitor of UBC9, fail to completely abolish the SUMO2/3 signal in the oocytes even after prolonged 48hrs incubation (Extended Data Fig. 8a-c); while 2) TAK-981, an inhibitor of SUMO-activating enzyme (SAE), achieved sufficient level of inhibition within few hours of treatment^48^ (Fig. 6a,b and Extended Data Fig. 8a,d). SUMO2/3 modification has been shown to be critical for the maintenance of centromeric cohesion during post-anaphase/telophase in oocytes^50^. Considering this, the oocytes were treated at the GV stage and the inhibitor carefully washed off before allowing the oocytes to progress through meiosis (Fig. 6a).

**Figure 6.**
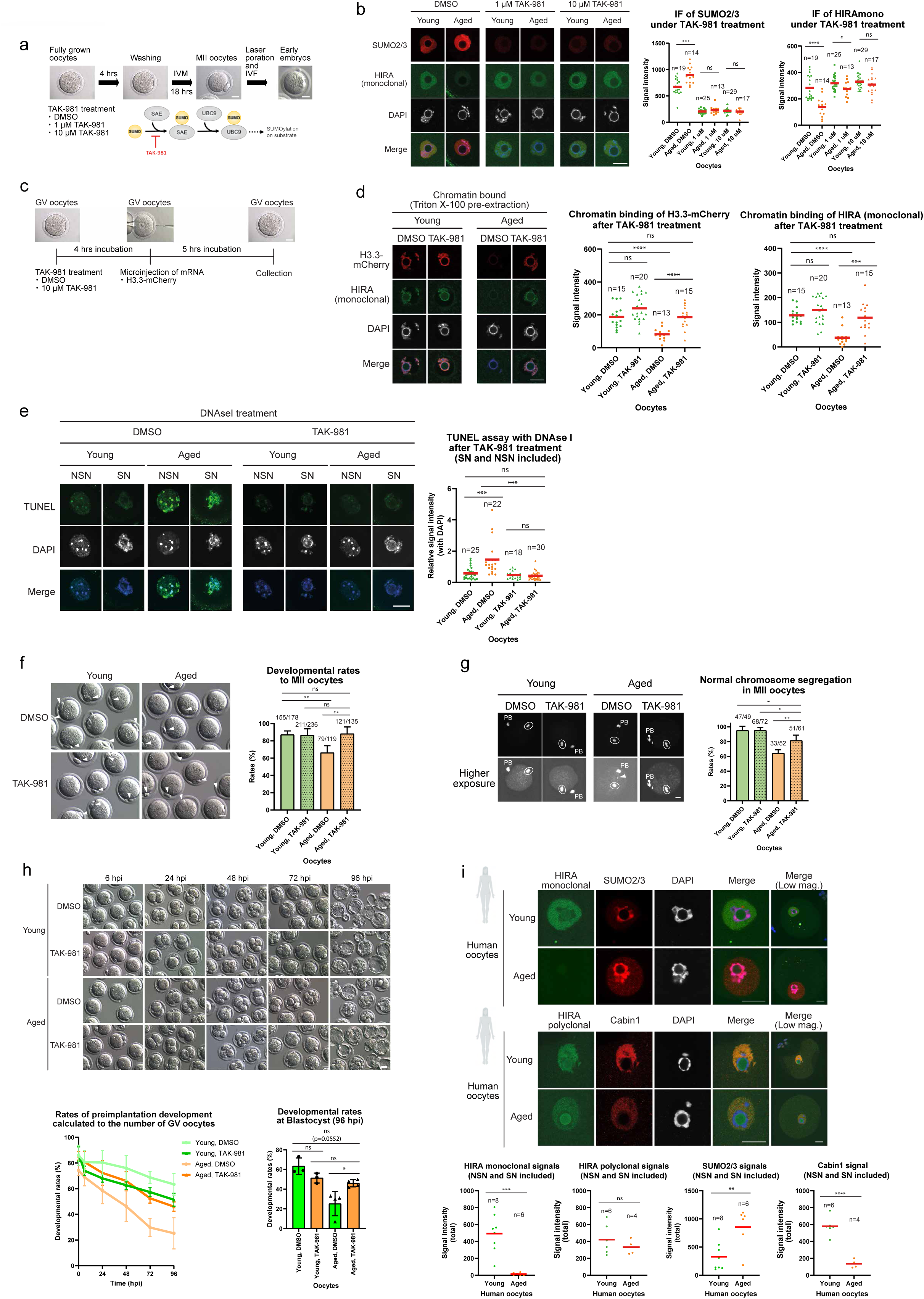
The effect of TAK-981 treatment on oocyte quality and evaluation of conservation in human oocytes. a. Experimental scheme. Fully grown mouse oocytes were collected for TAK-981 treatment. TAK-981 is an inhibitor of SUMO-activating enzyme (SAE). b. The efficiency of TAK-981 treatment and the effect on HIRA localisation. The DMSO treatment used as a control. n=number of analysed oocytes, ****p < 0.0001, ***p < 0.001, *p < 0.05, ns – not significant. Red bars indicate mean. Scale bar 20μm. c. Scheme of TAK-981 treatment followed by mRNA microinjection into GV oocytes. d. The effect of TAK-981 treatment on chromatin incorporation of H3.3-mCherry. The treatment of TAK-981 leads to increased chromatin incorporation of H3.3-mCherry in the aged oocytes, consistent with increased HIRA chromatin localisation. n=number of analysed oocytes, ***p < 0.001, ****p < 0.0001, ns – not significant. Red bars indicate mean. Scale bar 20μm. e. Z-Stacked images of DNAseI treatment followed by TUNEL assay in oocytes treated with 10μM TAK-981 (right) or DMSO (control, left). n=number of analysed oocytes, ***p < 0.001, ns – not significant. Red bars indicate mean. Scale bar 20μm. f. Efficiency of oocyte maturation to MII stage. In vitro maturation of DMSO (top) or TAK-981 (bottom) treated fully grown oocytes. Numbers above each bar indicate the number of MII oocytes over the total number of treated oocytes. Arrowheads show 1st polar body. Scale bar 30μm. **p < 0.01. ns - not significant. Bar shows SD. g. Chromosome condensation and segregation in MII oocytes treated with TAK-981. Higher exposure images shown to detect the segregation errors. Numbers above each bar indicate the number of MII oocytes over the total number of treated oocytes. Arrowheads show chromosome segregation error. Circles show mitotic chromosomes. PB, polar body. Scale bar 20μm. **p < 0.01, *p < 0.05, ns – not significant. Bar shows SD. h. Preimplantation development of embryos following TAK-981 treatment of oocytes. Scale bar 30μm. Efficiencies were calculated based on the number of oocytes isolated from the ovary. Significant differences were detected at blastocyst stages between “Aged, DMSO” and the others. *p < 0.05, ns – not significant. Bar shows SD. i. Localisation of HIRA (monoclonal antibody; top), SUMO2/3 (top), HIRA (polyclonal antibody; bottom), and Cabin1 (bottom) in human oocytes at SN stage. 4-8 oocytes were examined for per experimental group. Scale bar 20μm. n=number of analysed oocytes, **p < 0.01, ***p < 0.001, ****p < 0.0001, ns – not significant. Red bars indicate mean.

Although incubation with both 1 μM and 10 μM TAK-981 for 4hrs completely diminished SUMO2/3 signal in the oocytes’ nuclei (Fig. 6a,b), only 10 μM TAK-981 treatment was sufficient to fully recover signal for the anti-HIRA monoclonal antibody (Fig. 6b). This also abolished the association between HIRA and SUMO2/3 in the PLA assay (Extended Data Fig. 8d,e) and restored HIRA-Cabin1 interaction (Extended Data Fig. 8f). We hence decided to use 10 μM TAK-981 for all further experiments. This treatment regimen (Fig. 6a) restored the ability of aged oocytes to incorporate H3.3 (Fig. 6d), which resulted in decreased DNAseI sensitivity (Fig. 6e, Extended Data Fig. 8g), increased in vitro development rates to MII oocytes (89% with TAK-981 vs 66% with DMSO, Fig. 6f and Extended Data Fig. 9a) and significantly improved chromosome segregation defects in the aged oocytes (Fig. 6g and Extended Data Fig. 9b) thus leading to a substantial cumulative improvement of aged oocyte maturation (Extended Data Fig. 9c)

Finally, to confirm the improved quality of the treated aged oocytes, we decided to assess their developmental potential following in vitro fertilisation (IVF). Similar to the experiments using microinjection of HIRAmut (Fig. 5g), we switched to B6CBA F1 mouse strain, that showed the same restoration of aged oocyte quality in response to the TAK-981 treatment (Extended Data Fig. 9c). Preimplantation embryos derived from aged F1 oocytes were severely developmentally compromised in comparison to embryos generated using young oocytes (Fig. 6h). Intriguingly, the developmental competence was restored by TAK-981 treatment, as documented by the rate of development to the blastocyst stage (28% untreated vs 46% treated aged oocytes, in comparison to 53% of treated young controls, Fig. 6h, Extended Data Fig. 9d, Extended Data Table 3). Furthermore, upon transfer to pseudo-pregnant dams, the embryos that had originated from TAK-981 treated oocytes gave rise to healthy pups (Extended Data Fig. 9e) that showed normal fertility in adulthood (Extended Data Fig. 9e). These findings thus provide direct evidence that inhibition of SUMOylation restores efficient chromatin maintenance, which in turns leads to better maturation, improved chromosome segregation and increased developmental potential of the aged oocytes.

Finally, given the high sequence conservation (Extended Data Fig. 6b), we asked whether the observed phenomenon might be conserved in humans. Human young (21-32 years) and aged (35-44 years) GV oocytes were obtained from egg donors with no known history of infertility following hormonal stimulation. Immunofluorescence staining strikingly revealed reduced reactivity to HIRA monoclonal antibody as well as an increase in SUMO2/3 in the aged human oocytes. This was additionally accompanied by reduced Cabin1 signal in the nuclei of aged human GV oocytes, supporting the notion that the here uncovered age-related defect in the structural chromatin maintenance is conserved (Fig. 6i).

## Discussion

Age-related decline in female fertility has been directly linked to a progressive decrease in the quality of oocytes^52^. This has been previously attributed to several factors including: erosion of sister chromatid cohesion leading to frequent chromosome mis-segregation, changes in gene expression and epigenetic regulation^27,32^, altered mRNA translation and nucleophagy^53–55^, and also toxic effects of the somatic gonadal environment^56^ and diet^57^.

Here we show that aging mouse oocytes have severely reduced ability to incorporate H3.3 leading to their failure to maintain chromatin integrity (Fig. 2 and Extended Data Fig. 1). Our previous work uncovered that developing mouse oocytes critically rely on the histone chaperone HIRA and H3.3 incorporation to sustain normal transcriptional regulation, efficient chromosome segregation and high developmental potential^10^. We now show that in the aging oocytes, HIRA is modified by SUMO2/3, which leads to reduced interaction with CABIN1, reduced chromatin occupancy and attenuated H3.3 incorporation (Fig. 2 and Fig. 3). This in turn results in altered chromatin structure as shown by increased nuclease sensitivity and chromosome segregation errors (Fig. 1c, Fig. 2c,d and Extended Data Fig. 1) drawing clear mechanistic parallels between oocyte aging and the previously reported HIRA knockout phenotype^10^.

The HIRA histone chaperone complex mediates replication-independent deposition of the histone variant H3.3 into chromatin through core subunits, HIRA, UBN1, and CABIN1 next to transiently associating with histone chaperone ASF1a^39,46,58,59^. Notably, HIRA trimerization is critical for Cabin1 binding and its function^46^. Disruption of trimerization abolished CABIN1 binding entirely while UBN1 and ASF1a interactions are maintained, indicating that CABIN1 recruitment is exclusively dependent on the trimeric HIRA^46^. In agreement with this, our Alphafold predictions suggest that SUMOylation of HIRA could disrupt trimerization and also direct interface between HIRA and Cabin1 (Extended Data Fig. 5). Taken together, our results suggest that HIRA SUMOylation affects HIRA function by attenuating Cabin1 interaction. We also note that the recently reported cryo-EM structure of the human HIRA complex supports our findings underlying the importance of HIRA-Cabin1 interaction for chromatin association and function of the HIRA complex.

Critically, this newly uncovered molecular mechanism allowed us to design and test targeted interventions: we show that restoring the aging oocyte’s capacity to incorporate H3.3 by either providing non-SUMOylatable HIRA (Fig. 4, 5 and Extended Data Fig. 6, 7), or by small molecule inhibition of the SUMO pathway (Fig. 6a-h, Extended Data Fig. 8, 9) improves overall chromatin structure and alleviates both oocyte maturation and chromosome segregation defects (Fig. 5c-f, 6e-g, Extended Data Fig. 7, 8, 9a-c). This in turn results in improved developmental potential of the aged oocytes (Fig. 5g, 6h; Extended Data Fig. 9d-f and Extended Data Table 3).

We note, that although both approaches improved the oocyte quality, small molecule inhibition of SUMOylation had a superior effect on the quality of oocytes and the preimplantation development rates (Fig. 5e-g, Fig. 6f-h, Extended Data Fig. 7 and 9). We stress that these interventions generate mechanistically two distinct situations: Expression of mutated HIRA does not affect the existing pool of the endogenous SUMOylated molecules, which it has to outcompete to generate any effect. Very high level of HIRA expression is additionally toxic. To the contrary, SUMO pathway exists in the state of dynamic equilibrium, the use of TAK-981 disrupts this equilibrium leading to rapid SUMO removal from the existing modified proteins by the present SUMO proteases. The equilibrium in the pathway and the SUMO modifications can be subsequently restored following the removal of the inhibitor. This is particularly important given the reported role of SUMO in the final stages of oocyte maturation.

Our data show that the aging oocytes progressively accumulate SUMO2/3; our short TAK- 981 treatment thus provides a SUMO system “reset” allowing for deSUMOylation of HIRA and restoration of normal chromatin structure through renewed incorporation of H3.3 (Fig. 6 and Extended Data Figs.8,9). Critically, the SUMO pathway equilibrium is restored following the careful wash out of the drug with the subsequent 18hrs of IVM incubation allowing for the re-equilibration in the SUMO pathway (Fig. 6a).

While we could recapitulate the effect of SUMO inhibitor using microinjection of mutated HIRA suggesting that the effect on the HIRA itself drives the phenotype; we acknowledge that other components of the HIRA complex might also be subject to SUMO modification. In this context, both UBN1 and CABIN1 have been previously reported to undergo SUMOylation^60^ . The use of small molecule SUMO inhibitor would naturally impact those modifications as well.

Age related fertility decline is a gradual process. Our results show that by the week 40 the majority of HIRA molecules are postranslationally modified in the mouse oocytes, while the overall HIRA protein levels are still unchanged. This is then followed by a loss of HIRA protein by week 65-66, coinciding with the peak of SUMO2/3 accumulation, at the time when majority of C57Bl6 females are already infertile (Fig. 3d,e)^26^. Our observations thus raise important questions regarding the nature of upstream molecular triggers and why is HIRA modified so early in the aging process (before the SUMO2/3 reaches the maximum – Fig. 3d-f). To this point, both inflammation and tissue fibrosis have been reported to increase in aging ovary starting from between 7-9 months (30-40 weeks) of age and are abundant by 12 months of age. This is accompanied by increase in proinflammatory signalling and signs of tissue stress. Critically, SUMO2/3 has been linked to stress response and also reported to increase in direct response to interferon signalling and oxidative stress. Both increase in interferon signalling and oxidative stress are also well documented features of aging oocytes. The observed SUMO2/3 accumulation is thus likely reflecting both the changes in surrounding ovarian milieu (increase in ovarian tissue inflammation and fibrosis), as well as upregulated stress response and changes in proteostasis that accompany oocyte aging^9^

Furthermore, HIRA itself has been reported to associate with PML bodies in response to interferon signalling. We also note that HIRA has been reported to be SUMOylated upon heat shock and proteostatic stress in cultured cells showing one of the highest stress responsiveness. These facts not only cumulatively corroborate our findings of HIRA being modified early in the oocyte aging process, but critically, point towards the existence of a distinct time window for a possible intervention.

Although our mechanistic work has been carried out using aged mouse oocytes, we show that the aged human oocytes recapitulate our fundamental findings from the mouse (Fig. 6i). We observed loss of reactivity to HIRA monoclonal antibody and increased SUMO2/3, while the overall signal for HIRA (polyclonal antibody) remains constant. We have also observed reduced signal for Cabin1, collectively pointing towards altered functionality of the HIRA complex in aging human oocytes. In addition, HIRA locus has been linked with the premature ovarian failure in humans in a recent GWAS study^61^, strengthening the link between HIRA function and oocyte health, underlying the importance of our work and opening the route towards possible therapeutic interventions.

Finally, loss of epigenetic information has been identified as a cause of mammalian aging, with the described phenomenon attributed primarily to the accumulation of DNA damage^62^. Here we show that in the oocyte, aging is accompanied by a loss of ability to maintain normal chromatin structure with wide ranging implications for maintenance of genome integrity and epigenome stability. Our work has thus uncovered a novel mechanism underpinning age related loss of epigenetic identity and cellular functions with possible implications for other long lived postmitotic cells.

## Methods

### Mice

All animal experiments were approved by UK Home office under Project License PPL PP1838178, and followed all relevant guidelines and regulations. Mouse rooms have a 12-hour light–dark cycle and the room temperature was maintained at 20–24 °C. The relative humidity was kept at 45–65%.

### Collection of GV Oocytes, in vitro Maturation (IVM), its in vitro Fertilisation (IVF), Embryo Culture, and Embryo Transfer into Recipient Females

Fully grown GV oocytes were collected from C57BL/6J strain female mice; young oocytes from 6-10 weeks old females, aged oocytes from 40-104 weeks old females. Collection of spermatozoa, oocytes at GV and MII stages generated using IVM, and fertilised embryos was performed as described^63,64^. For IVM, GV oocytes were subjected to a Hepes-buffered 1:1 mixture of TYH and αMEM (TαM) (gibco). For IVM-IVF, all oocytes were prepared by IVM. A small incision was made in the zona pellucida using laser to facilitate sperm penetration by NaviLase (Octax) and the sperm suspension was capacitated in human tubal fluid (HTF) medium for 1.5 h at 37 °C under 5% CO2 in humidified air. Spermatozoa were collected from the cauda epididymis of C57BL/6J male mice. Fertilized embryos were cultured in potassium simplex optimized medium (KSOM)^65^ at 37 °C under 5% CO2 in humidified air. The embryos were assessed by both timing of the progression (ie assessment at particular hours post insemination (hpi), 24 hpi: 2 –cell stage, 48 hpi: 4-8 cell stage, 72 hpi: Morula stage, and 96 hpi: Blastocyst stage) and embryo morphology. 2C stage – presence of two equal size blastomeres, 4-8C stage - presence of 4-8 equal cells, Morula stage – embryos consisting of a compacted mass of approximately 8–16 blastomeres, with no visible blastocoel cavity, blastocyst – judged by the emergence of a fluid-filled cavity known as the blastocoel. To obtain pups, B6CBA F1 strain females were used. The embryos at blastocyst stage were transferred to into recipient females, and the females were subjected to cesarean section at E19.5, and the pups were collected.

### Single oocyte RNA sequencing analysis of young and aged oocytes

Five mice were used for the oocyte collection. The pooled oocytes with a diameter of more than 60 μm were selected by using a hold of micromanipulator with the size adjusted. This was performed as described^10^. In brief, cDNA synthesis and amplification was performed directly on single young and aged oocytes with the SMART-Seq® v4 Ultra® Low Input RNA Kit (Clontech). 0.5 μl of ERCC RNA spike-in mix 1 (Life Technologies), diluted 1:10^5^, was added to each reaction prior to cDNA synthesis. The amplified cDNA was fragmented by using a Covaris S2 instrument (Covaris) and converted to sequencing libraries following the NEBNext® Ultra™ II DNA Library Prep (NEB) using the NEBNext Multiplex Oligos for Illumina (NEB).

Reads adapter trimmed using trimmomatic (v 0.39) and were mapped against GRCm38 (mm10) and the ERCC spike sequences using STAR (v 2.7.7a)^66^, which was also used to prepare gene-level counts. Additionally, transposable element (TE) counts were prepared using TEtranscripts (v 2.2.3)^67^, summarising counts to the family level. Generated count data were normalised and differentially expression assessment performed using DESeq2^68^. Differentially expressed genes and TEs were defined with Benjamini-Hochberg adjusted P < 0.05 using the Wald test. Gene Set Enrichment Analysis was performed using GSEA software^69,70^ using the ranks generated by DESeq2.

### Preparation of human oocytes

Human oocytes were obtained by Newcastle Fertility Centre (Newcastle upon Tyne, UK) and experiments conducted under an HFEA- research license (R0152) with Health Research Authority approval from Newcastle and North Tyneside Research Ethics Committee. Informed consent was obtained from all donors by research nurses who were not directly involved in the research, or in the provision of clinical treatment. Immature oocytes donated specifically for research (n = 40 oocytes from 26 donors, age range 21 - 44 years) were used. Donors were stimulated with a fixed dose antagonist protocol. They were commenced on recombinant human follitropin alfa injections (225 IU subcutaneous daily for 7-12 days) for ovarian stimulation. GnRH antagonist, Cetrorelix or Ganirelix (0.25 mg subcutaneous daily) was added five days after commencement of the stimulation regime to prevent premature LH surge and continued till the trigger day. The follicular development was monitored by ultrasound scan to decide the day of trigger. The ovulation trigger used was either recombinant hCG (0.25 mg) or GnRH agonist, Buserelin (0.5 mg). The oocytes were collected at 21.5– 40 hours post-trigger by transvaginal ultrasound-guided follicle aspiration. The surrounding cumulus cells were removed with HYASE (Vitrolife, Sweden) diluted in G-MOPS PLUS (Vitrolife, Sweden). Oocytes were vitrified using RapidWarm oocyte kit (Vitrolife, Sweden) if they were not used immediately. They were stored in liquid nitrogen until required. Oocyte warming was performed using RapidWarm oocyte kit (Vitrolife, Sweden). In accordance with HFEA Directions (Human Fertilisation and Embryology Authority, 2018), egg donors received financial compensation of £500 per donation cycle as approved by ethics committee and HFEA. Immature oocytes (n = 14 from 4 donors, age range 32-37 years) were donated for research by couples undergoing standard infertility treatment.

### Immunocytochemistry and Microscopy

Mouse oocytes were fixed in 4% paraformaldehyde (PFA) for 15min at room temperature. Human oocytes were fixed in 4% PFA at 4 °C for 1 hour. Immunocytochemistry was performed as described^63^. After washing with PBS containing 1% bovine serum albumin (BSA), the fixed oocytes were then treated with PBS containing 0.5% Triton X-100 at RT for 40 min. For pre-extraction method, oocytes were treated with 0.2% Triton X-100 (SIGMA) in PBS for 30 s before the fixation. They were then incubated with the primary antibodies (Extended Data Table 4) in PBS containing 1% BSA at 4 °C overnight. After washing, they were reacted with the secondary antibodies as appropriate (Extended Data Table 4) at RT for 1 h. Specimens were mounted on glass slides in Vectashield mounting medium with DAPI (Vector Laboratories). HIRA knockout ES cells were used for validation of the HIRA monoclonal and polyclonal antibodies by IF staining. The ES cells were fixed in 4% PFA for 20 min at room temperature. After washing, the fixed cells were then permeabilised with PBS containing 1% BSA and 0.1% Triton X-100 at RT for 30 min. The primary and secondary antibody reaction was treated with the permeabilised solution. Finally, the Specimens were imaged using a Leica TCS SP5 confocal microscope with a 40x objective and Z-step size of 0.5 μm in the case of Z stack scanning. ImageJ software (NIH, Bethesda, MD, USA; http://rsbweb.nih.gov/ij/) was used to quantify DAPI staining and antibody signals for total signal intensity. At least three independent replicates were performed for each experiment.

### Detection of newly synthesized RNA by EU incorporation

Click-iT RNA imaging kit (Invitrogen) was used for detection of newly synthesized RNA by EU incorporation. Briefly, 1mM of EU labelling was added to culture medium and oocytes were cultured for 1 hour. The oocytes were then fixed in 4% PFA, followed by a treatment with 0.5% TritonX-100 in PBS for 15 min. They were then incubated with Click-iT reaction cocktail for 30 min at room temperature. Specimens were mounted on glass slides in Vectashield mounting medium with DAPI and observed as described above.

### Proximity Ligation Assay (PLA)

PLA was performed by using Duolink® In Situ PLA kit (SIGMA) according to the manufacturer’s instructions. In brief, the fixed oocytes were permeabilised as described above. After washing in PBS containing 30 mg/ml bovine serum albumin, the oocytes were incubated with Duolink® Blocking Solution at 37 °C for 1 h and then incubated with the primary antibodies (Extended Data Table 4) in Duolink® Antibody Diluent at 4 °C overnight. Washing steps were carried out using washing buffer A or B provided in the kit. After washing, probe incubation was performed using probe solutions in Duolink® Antibody Diluent at 37 °C for 1 h followed by the incubation with ligation solution at 37 °C for 30 min. After washing, amplification reaction was performed using polymerase in a 1x amplification buffer according to the manufacturer’s instructions. After additional washing, the oocytes were mounted on glass slides in Vectashield mounting medium with DAPI (Vector Laboratories) and imaged as described above.

### TUNEL assay for detection of the DNAse I sensitivity

TUNEL assay was performed using Click-iT TUNEL Alexa Fluor Imaging Assay (Life Technologies) according to manufacturer’s instructions. For DNAse I treatment, the oocytes were incubated with DNAse I (Roche) at 0.1 U/μl or at various concentrations for 30 min at 37 °C. At least three independent replicates were performed for each experiment.

### NicE-view labelling of accessible chromatin in oocytes\

Fixed oocytes were permeabilised for 40 min at room temperature in permeabilization solution (1% BSA-PBS + 0.5% Triton X-100), washed in 1% BSA-PBS, flash-washed once in PBS(−), and equilibrated for 30 min at room temperature in 1× NEBuffer 2 (NEB, B7002S) diluted in nuclease-free water. Accessible chromatin labelling buffer (Extended Data Table 5) was prepared immediately before use at 50 µl per reaction. Oocytes were flash-washed in 1× NEBuffer 2, transferred into 50 µl of labelling buffer and incubated for 2 h at 37 °C in the dark. Reactions were stopped by adding 0.5 µl EDTA and 0.2 µl RNase A directly to the well under a dissecting microscope and incubating for a further 20 min at 37 °C. The reaction mixture was removed by mouth pipette and replaced with 200 µl of 1% BSA-PBS pre- warmed to 55 °C. Oocytes were incubated for 30 min and washed twice more in 1% BSA- PBS for 30 min per wash. Oocytes were mounted on glass slides with spacers in VECTASHIELD antifade mounting medium containing DAPI and imaged by confocal microscopy.

### In Vitro mRNA Synthesis and microinjection

This was performed as described^71^. In brief, each expression plasmid of H3.1-GFP, H3.2- GFP, and H3.3-GFP were prepared previously^10^. Each cassette of EGFP-HIRAwt, EGFP- HIRAmuts, and H3.3-mCherry was generated by gene synthesis (GeneCust, France) and cloned into pCDNA3.1 Myc HisA by the GeneCust. The prediction of SUMOylation sites on HIRA was performed by GPS-SUMO 2.0 (https://sumo.biocuckoo.cn/). The pcDNA-HIRA- polyA (Addgene #59781) and pGEMHE-NLS-mEGFP (Addgene #105527) were also used for in vitro mRNA synthesis. Amplification and poly(A) tailing of these mRNAs were performed using mMESSAGE mMACHINE T7 Ultra (Ambion) for each construct. GV oocytes were injected with the mRNAs using a Piezo-driven micromanipulator and then the oocytes were transferred to HTF containing 203.5 μM dbcAMP (SIGMA) and 100 μM IBMX (SIGMA) until collection.

### Production of ES Cells with stably expressed SUMO2Q87R and Purification of Flag- HIRA Protein

The E14 mESCs stably expressing SUMO2Q87R were generated by transfection of the pcDNA3.1-StrepHA-SUMO2Q87R (Addgene #66868) using Lipofectamine™ 3000 (Invitrogen), followed by geneticin (Gibco) treatment for drug-selection. The geneticin-resistant cells were confirmed by the protein expression and its localisation by western blot and Immunostaining. The cassette of Flag-HIRA-His was subcloned into pCAGIG plasmid (Addgene #11159) to induce the transient expression in the ES cells. The transfection was performed using Lipofectamine™ 3000. Cells were treated with 10 μM MG-132 (SIGMA) for 7 hours prior to harvesting. At 48 h post-transfection, the cells were harvested and lysed with Lysis buffer containing 20 mM Tris-HCl, 0.5% NP-40, 150 mM NaCl, and 1x protease inhibitor (Roche). The lysate was sonicated by Bioruptor® (Diagenode) (30 sec ON/ OFF, 5 cycles, low intensity) at 4 °C. After centrifugation at 12,000 g for 10 min, the supernatants were incubated with Anti-FLAG® M2 Magnetic Beads (SIGMA) at 4 °C overnight with gentle rotation. The magnetic beads were washed 5 times with Wash buffer containing 50 mM HEPES pH7.4, 1 mM EDTA, 150 mM NaCl, and 1x protease inhibitor. The elution buffer containing 50 mM HEPES pH7.4, 1 mM EDTA, 100 mM NaCl, and 300 ng/ μl of Flag peptide (SIGMA) was added to the beads and then the suspension was incubated for 30 min at 4 °C with rotation. The supernatant was used for WB analysis and MS analysis. The SUMO2Q87R was purified by Monoclonal Anti-HA Agarose (Millipore) with the same buffers and procedures as the Flag-HIRA purification.

### Western Blot analysis

The procedures were performed as described^63^. In brief, the proteins were resolved in 4– 20% Mini-PROTEAN® TGX™ Precast Gel (Bio-Rad) and transferred electrophoretically to polyvinylidene difluoride (PVDF) membranes (Bio-Rad). The membranes were blocked in 2% skimmed Milk (Marvel) at room temperature (RT) for 1 h, washed with phosphate-buffered saline containing 0.2% Tween 20 (PBST) and incubated at 4 °C overnight with relevant primary antibodies (Extended Data Table 4). The membranes were then washed in PBST, incubated with HRP-conjugated secondary antibodies at RT for 1 h, washed with PBST, and developed using Immobilon Crescendo Western HRP substrate (Millipore). The images were acquired using Amersham Imager 680 (GE healthcare).

### *in vitro* SUMOylation assay

In vitro SUMOylation reaction was performed using SUMOylation Kit (Enzo) according to the manufacturer’s instructions. The purified Flag-HIRA was reacted with 1x SUMO E1, 1xSUMO E2, 1x SUMO proteins, 1x Mg-ATP solution, and 1x SUMOylation Buffer at 37 °C for 1 h. The reaction was stopped by adding Laemmli SDS-sample buffer and used for WB analysis.

### SUMOylation inhibitor treatment

GV oocytes were treated with TAK-981 (Selleck) for 4 h and 2-D08 (SIGMA) for 48 h, respectively, to assess the efficiency of the SUMOylation inhibitors. Treatment with 10 µM TAK-981 was carried out for 4 hours to obtain sufficient efficiency.

### Structural analysis of the HIRA complex

Using the AlphaFold3 web server^73^, we predicted residues 1-705 of Cabin1 with one copy of ASF1a, H3C, H4, UBN1_1-200_ (residues 1-200) and trimeric full-length HIRA (Extended Data Fig. 5a-c). HIRA forms a trimer with 3-fold rotational symmetry, with Cabin1 binding on one side of the complex, bridging two HIRA monomers. Cabin1 “head” region, residues 32-230 interact with residues 393-406 and 950-1014 of one HIRA and Cabin1 residue 210-225 interacting with 862-962 of the adjacent HIRA. This is consistent with studies demonstrating HIRA trimerization is needed for Cabin1 binding, while HIRA residues 873–904 promote Cabin1 binding, but are not needed for trimerization^46^. ASF1a, H3C, H4 and UBN1_1-200_ (residues 1-200) form a complex which binds to the N-terminal WD40 domain of HIRA, on the opposite side of the HIRA trimer to Cabin1. This prediction is in agreement with previously published structure of ASF1a with the HIRA B-motif^74^ and ASF1a with UBN1_122-148_ (residues 122-148), H3C and H4^58^. The prediction confidence of HIRA was improved in trimeric form, with higher pLDDT score and PAE contacts where HIRA 450-462 associated with ASF1a (arrow in Extended Data Fig. 5b).

We mapped the putative SUMOylation site of HIRA (Extended Data Fig. 5a, in red). No SUMOylation sites are found in the interface with ASF1a, UBN1, H3C and H4, consistent with experimental results indicating these are still able to bind when HIRA is SUMOylated. However, the model suggests SUMOylation could interfere with the HIRA/Cabin1 interaction in two ways (Extended Data Fig. 5d, insert). Firstly, residue HIRA K1015 lies near the HIRA/Cabin1 interface, hence SUMOylation may disrupt this interaction. Secondly, residue K807, lies at the trimerization interface of HIRA, hence, SUMOylation of this site could disrupt trimerization, which is necessary for Cabin1 binding.

We then modelled HIRA hexamers, composed of two full length HIRA molecules and four truncated spanning residues 675-1015, with two copies of Cabin1_1-705_ (Extended Data Fig. 5e-g). The Cabin1 “head” region, residues 32-230 bound one HIRA trimer, while the “shoulder” region, residues 270-700 bound the other trimer, such that Cabin1, bridged the trimers, consistent experimental results for yeast and human complex^46,75,76^. Binding of the Cabin1 head region was consistent with that in the trimer model. Mapping the SUMOylation sites, we found HIRA K1015 lies near the HIRA/Cabin1 interface, while residue K807, lies at the trimerization interface of HIRA (Extended Data Fig. 5h). Additionally, we found residue K858 lies near the interface of the Cabin1 shoulder with HIRA (Extended Data Fig. 5i), which could further disrupt the Cabin1/HIRA interaction.

In conclusion, structural analysis suggests two mechanisms of how SUMOylation of HIRA could reduce Cabin1 association, either via direct perturbation of the interface, or via interfering with HIRA multimer structure.

### Statistical Analysis

For statistical analysis, we used GraphPad PRISM and performed unpaired t tests for the signal intensities. P values were calculated by two-tailed unpaired t tests. P values < 0.05 were considered significant. For the analysis of differentially expressed genes and TEs in scRNASeq were described above.

## Data availability

scRNA-seq data have been deposited in Gene Expression Omnibus (GEO) under GSE247932. Source data are provided with this paper.

## Acknowledgements

We thank D. Dormann and C. Whilding for help with microscopy and immunofluorescence data analysis; L. Game for help with next-generation sequencing; M. Sayell and G. Zimmerman for mouse husbandry; and the members of the Hajkova laboratory for discussions and revisions of the manuscript; Work in the Hajkova laboratory is supported by MRC funding (MC_US_A652_5PY70) and the ERC grant (ERC-CoG-648879–dynamic modifications) to P.H. Human oocyte work is supported by grants from Horizon 2020 (GermAge, 634113). Y.H. is a recipient of the Japanese Society for the Promotion of Science (JSPS) Postdoctoral Fellowship (PD) No. 201700058. Y.H. is supported by JSPS KAKENHI grant number 23K20043 (Dr. Masahito Ikawa).

## Author contributions

Y.H. and P.H. conceived the study. Y.H. performed the experiments and Y.H. and P.H. analysed the data. Y.H. and B.N. performed the scRNASeq and M.D. and G.Y. analysed the next-generation sequencing data. M.H. performed the NicE-view assay with the help of P.O.E. and S.P. Y.T. and M.H. performed the experiments using human oocytes. E.C. carried out the structural analysis of the HIRA complex. Y.H. and P.H. wrote the manuscript.

## Competing interests

Y.H and P.H are inventors on the related patent application GB2622495.6

## Figure Legends

**Extended Data Figure 1.**
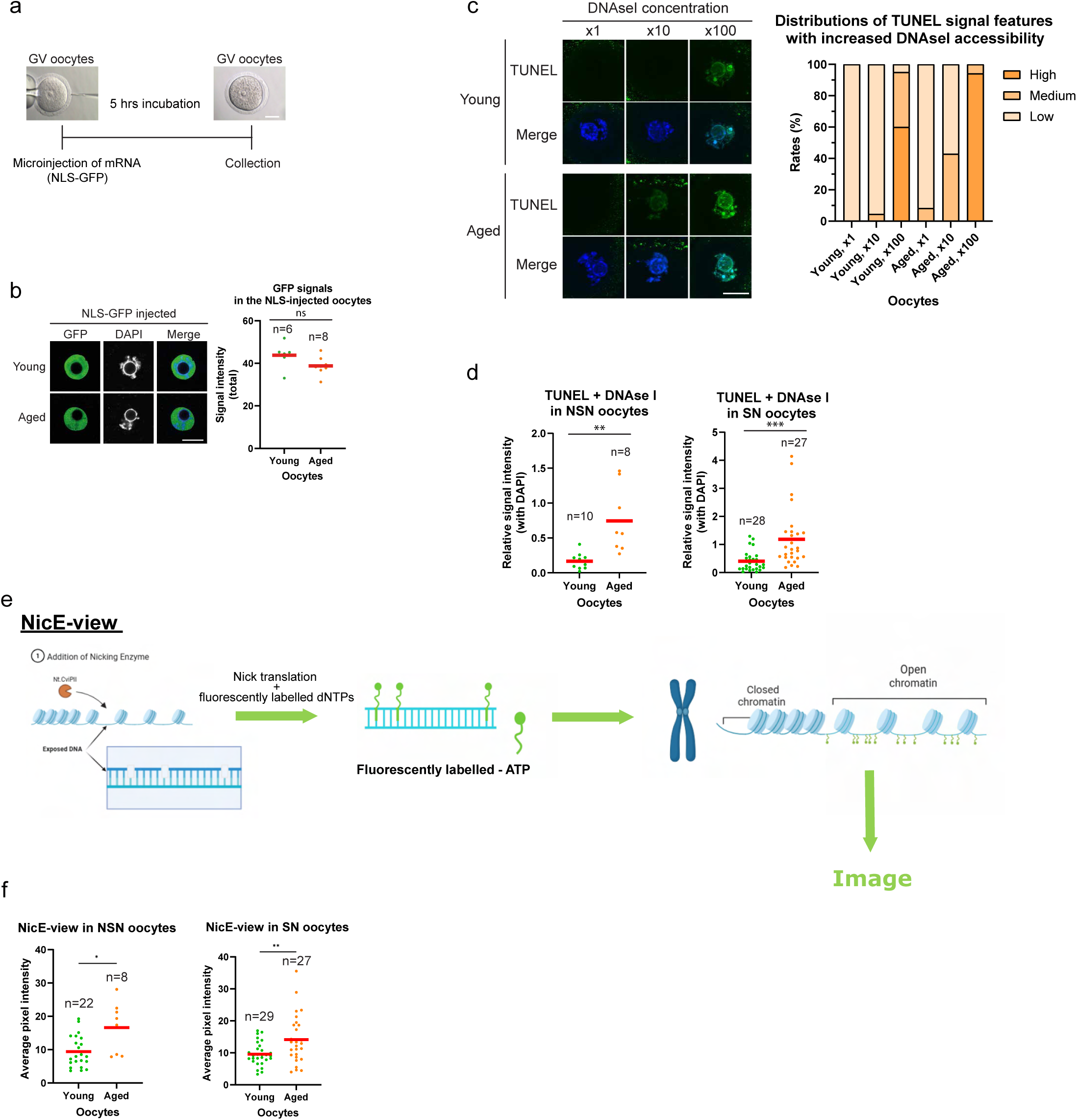
Chromatin dynamics is altered during aging. a. Scheme of microinjection of NLS-GFP mRNA into GV oocytes. b. Injection of NLS-GFP reveals no difference in translation efficiency or nuclear import between young and aged oocytes. n=number of analysed oocytes, ns - not significant. Red bars indicate mean. Scale bar 20μm. c. Z-Stacked images of treatment with different concentrations of DNAseI followed by TUNEL assay (left). The bar graph shows frequencies of signal intensities (right). Scale bar 20μm. d. Quantification of the TUNEL assay at NSN and SN stages, related to Figure 2c. n=number of analysed oocytes,**p < 0.01. ***p < 0.001. Red bars indicate mean. e. Schematic diagram of NicE-view assay. f. Quantification of the NicE-view assay at NSN and SN stages, related to Figure 2d. n=number of analysed oocytes, *p < 0.05. **p < 0.01. Red bars indicate mean.

**Extended Data Figure 2.**
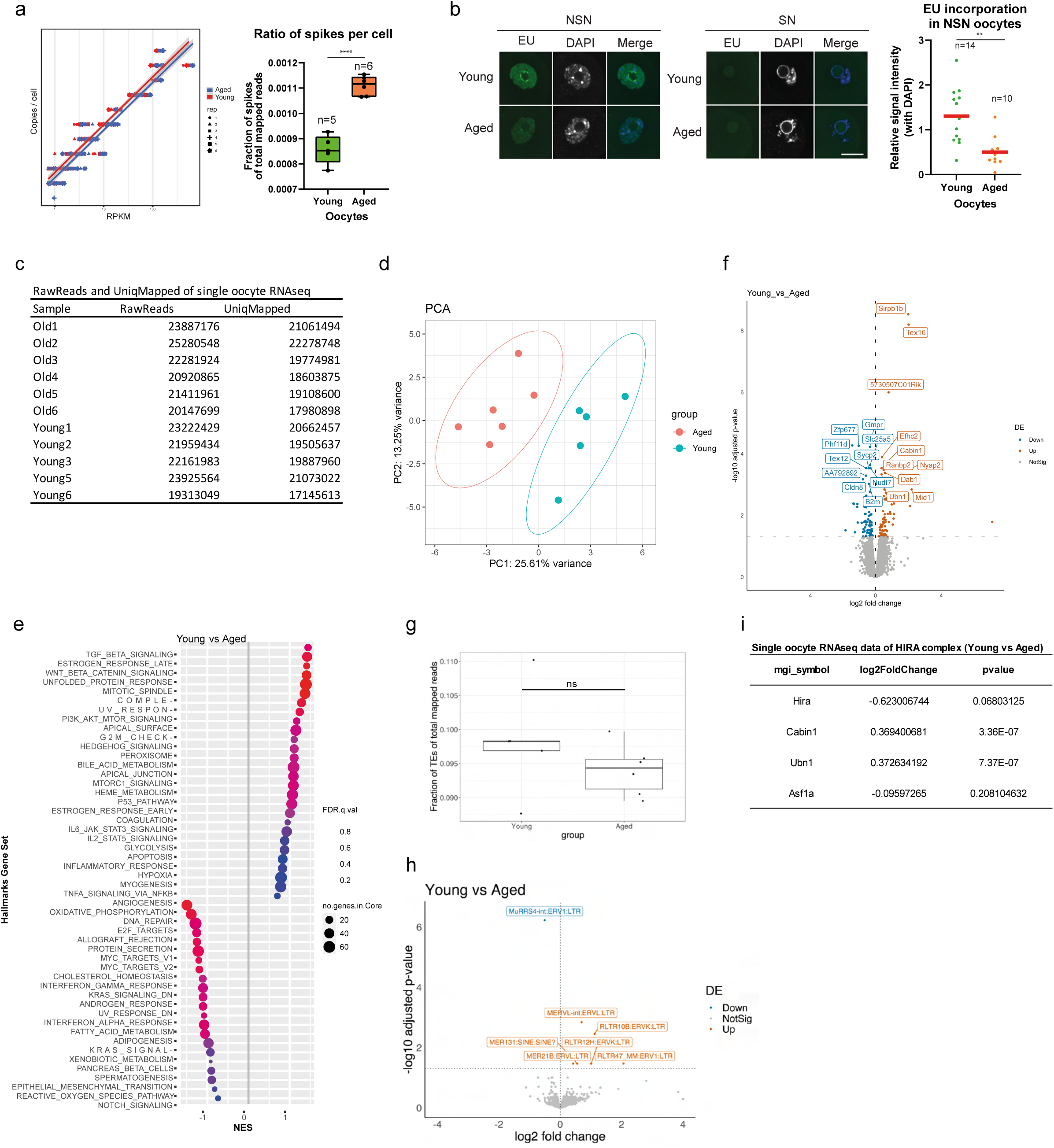
HIRA function is altered in aged oocytes. a. Comparison between obtained FPKM values and absolute copy number per cell for ERCC spike-in RNA in young and aged oocytes showing higher amplification in the latter. ****p < 0.0001, n = number of analysed single oocyte RNA Seq datasets. Bar shows SD. b. Detection of newly synthesised RNA by EU incorporation in young and aged oocytes at NSN and SN stages. Scale bar 20μm. Red bars indicate mean. n=number of analysed oocytes, **p < 0.01. c. Numbers of RawReads and uniqMapped reads in each of the single oocyte RNA Seq sample. d. PCA analysis of the single oocyte RNASeq datasets (young vs aged oocytes). e. GSEA between young and aged oocytes. f. Volcano plot depicting genes differentially expressed between young and old oocytes. Top 10 downregulated (blue) and upregulated (orange) genes based on the adjusted p-value are shown. g. TE fraction in total counts (single oocyte RNASeq).ns – not significant. h. Volcano plot depicting TEs differentially expressed between young and aged oocytes. Downregulated (blue) and upregulated (orange) TEs (young vs aged) based on adjusted p-value. i. Expression of selected components of the HIRA complex as detected in our single oocyte RNA Seq (Fold Change and statistical significance).

**Extended Data Figure 3.**
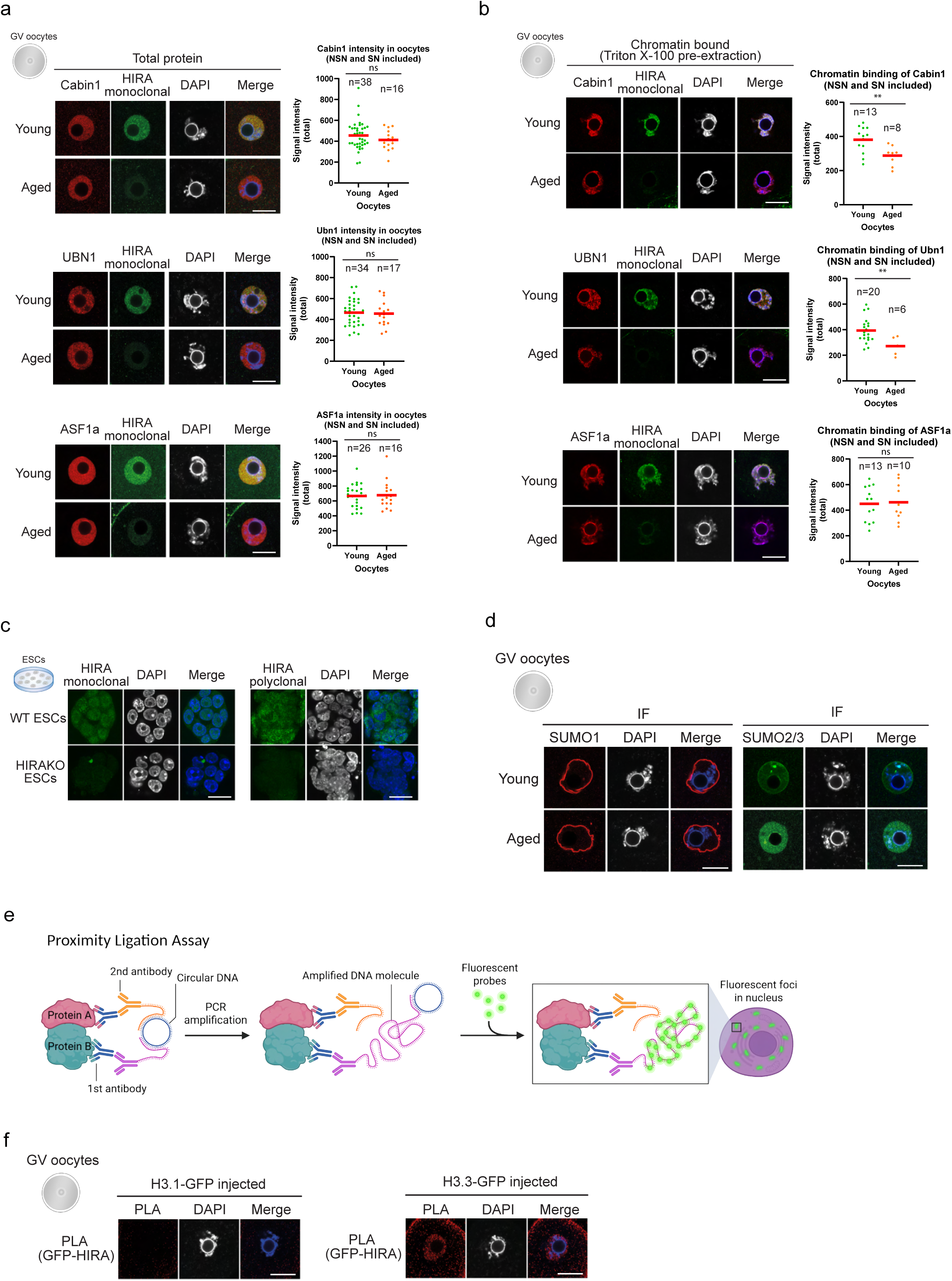
Characterisation of HIRA complex and SUMO proteins in oocytes. a. Localisation of HIRA, Cabin1, UBN1, and ASF1a in SN oocytes. No significant differences were detected. n=number of analysed oocytes. ns – not significant. Red bars indicate mean. Scale bar 20μm. b. Chromatin binding of HIRA, Cabin1, UBN1, and ASF1a in SN oocytes. n=number of analysed oocytes, **p < 0.01, ns – not significant. Red bars indicate mean. Scale bar 20μm. c. HIRA monoclonal and polyclonal antibodies were validated by using wt and HIRA KO mES cells. Scale bar 20μm. d. Nuclear membrane localisation of SUMO1 and nuclear localisation of SUMO2/3 at SN stage oocytes. Scale bar 20μm. e. Schematic diagram of PLA assay. Created with BioRender.com. f. Specificity of PLA assay was confirmed using microinjection of H3.1-GFP and H3.3- GFP, respectively, into GV oocytes followed by PLA using anti-GFP and anti-HIRA antibodies. The signal was detected only between H3.3 and HIRA, as expected. Scale bar 20μm.

**Extended Data Figure 4.**
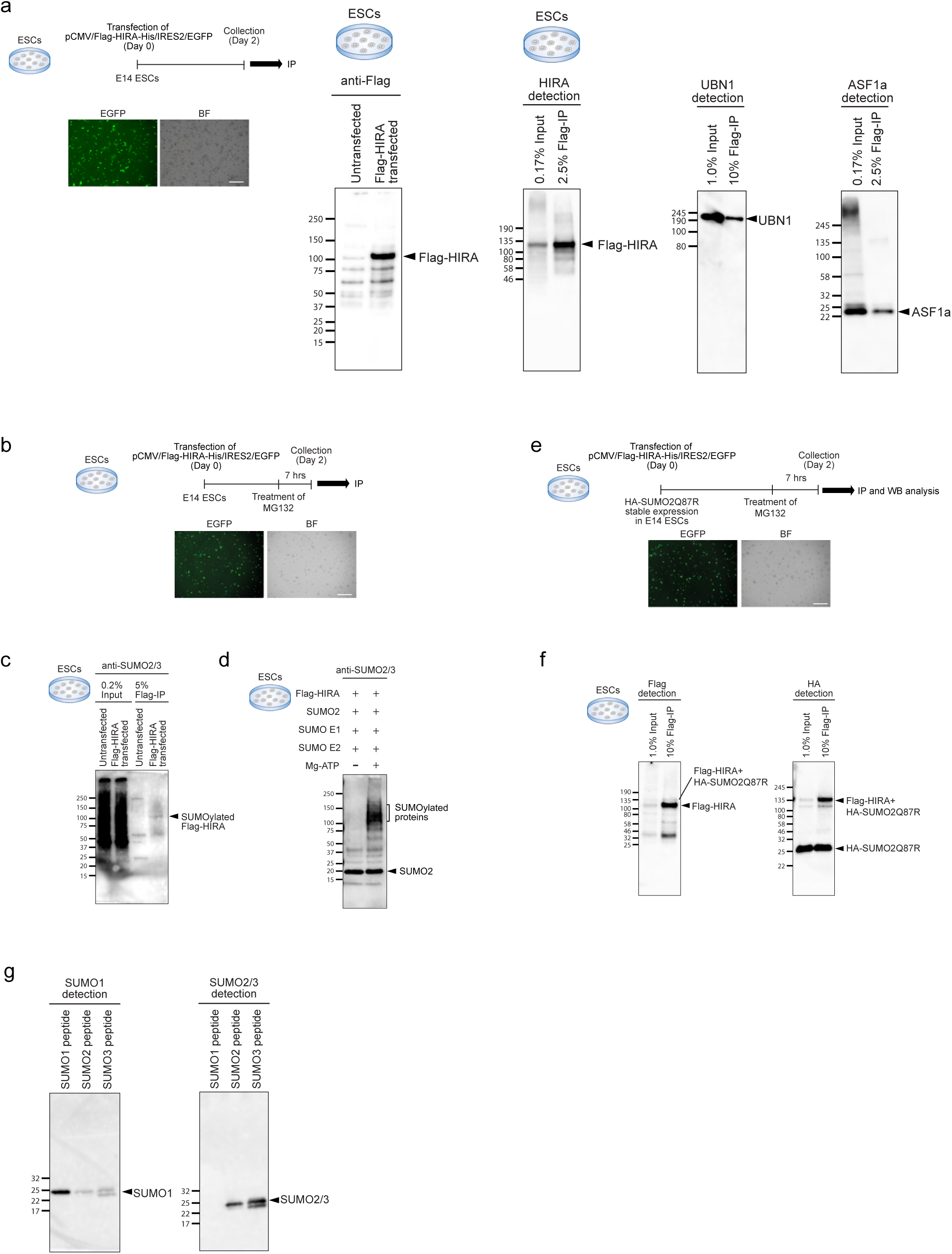
HIRA can be modified by SUMO2/3 in vitro and in cultured cells. a. Scheme for over-expression and purification (IP) of Flag-HIRA in E14 mESCs (left). BF, Bright Field. Scale bar, 500μm. Western blot analysis to confirm Flag IP efficiency by using anti-Flag and anti-HIRA antibodies (middle). HIRA interacting partners, UBN1 and ASF1a, were also detected (right). b. Scheme for over-expression of Flag-HIRA in E14 mESCs followed by MG132 treatment to avoid degradation of SUMOylated proteins. Scale bar 500μm. c. SUMOylated HIRA could be detected in the IP fraction by using anti-SUMO2/3 antibody. d. *In vitro* SUMOylation assay using Flag-HIRA as a substrate. e. mESCs stable expressing HA-SUMO2Q87R were transfected with Flag-HIRA followed by IP and WB analysis. Scale bar 500μm. f. Western blot analysis of Flag-HIRA IP shows the presence of SUMOylated HIRA using anti-Flag and anti-HA antibodies. g. The specificity of SUMO1 and SUMO2/3 antibodies as assessed by cross-reactivity in western blot.

**Extended Data Figure 5.**
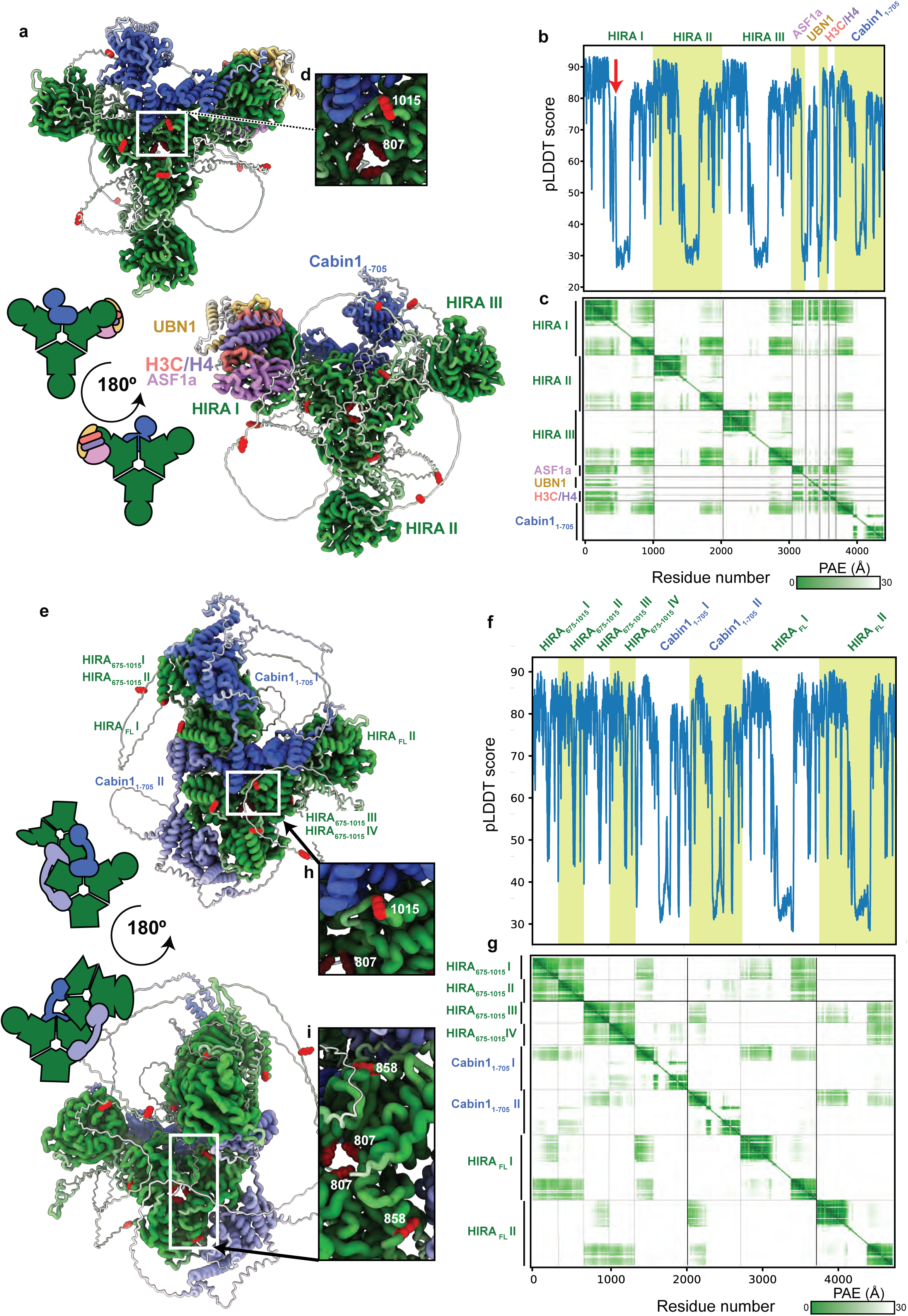
Structural analysis of the HIRA complex using Alphafold. For Full description see Materials and Methods. a. AlphaFold3 prediction of mouse HIRA complex. Putative HIRA SUMOylation sites are shown in red. b. pLDDT score denotes the prediction confidence of HIRA in trimeric form. Arrow showed association of HIRA 450-462 with ASF1a. c. PAE values are displayed on a green colour scale ranging from 0 to 30 Å, with dark green indicating low PAE. d. The model suggests SUMOylation could interfere with the HIRA/Cabin1 interaction. The putative SUMOylation sites of HIRA shown in red. e. Prediction of HIRA hexamers, composed of two full length and four truncated (spanning residues 675-1015) HIRA molecules, with two copies of Cabin1_1-705_. f. pLDDT score of HIRA hexamers and two copies of Cabin1_1-705_. g. PAE values are displayed on a green colour scale ranging from 0 to 30 Å, with dark green indicating low PAE. h. The SUMOylation sites at K807 and K1015 lies at the trimerization interface of HIRA and near the HIRA/Cabin1 interface, respectively. The putative SUMOylation sites of HIRA shown in red. i. Additional SUMOylation sites which could interfere with the HIRA/Cabin1 interaction.

**Extended Data Figure 6.**
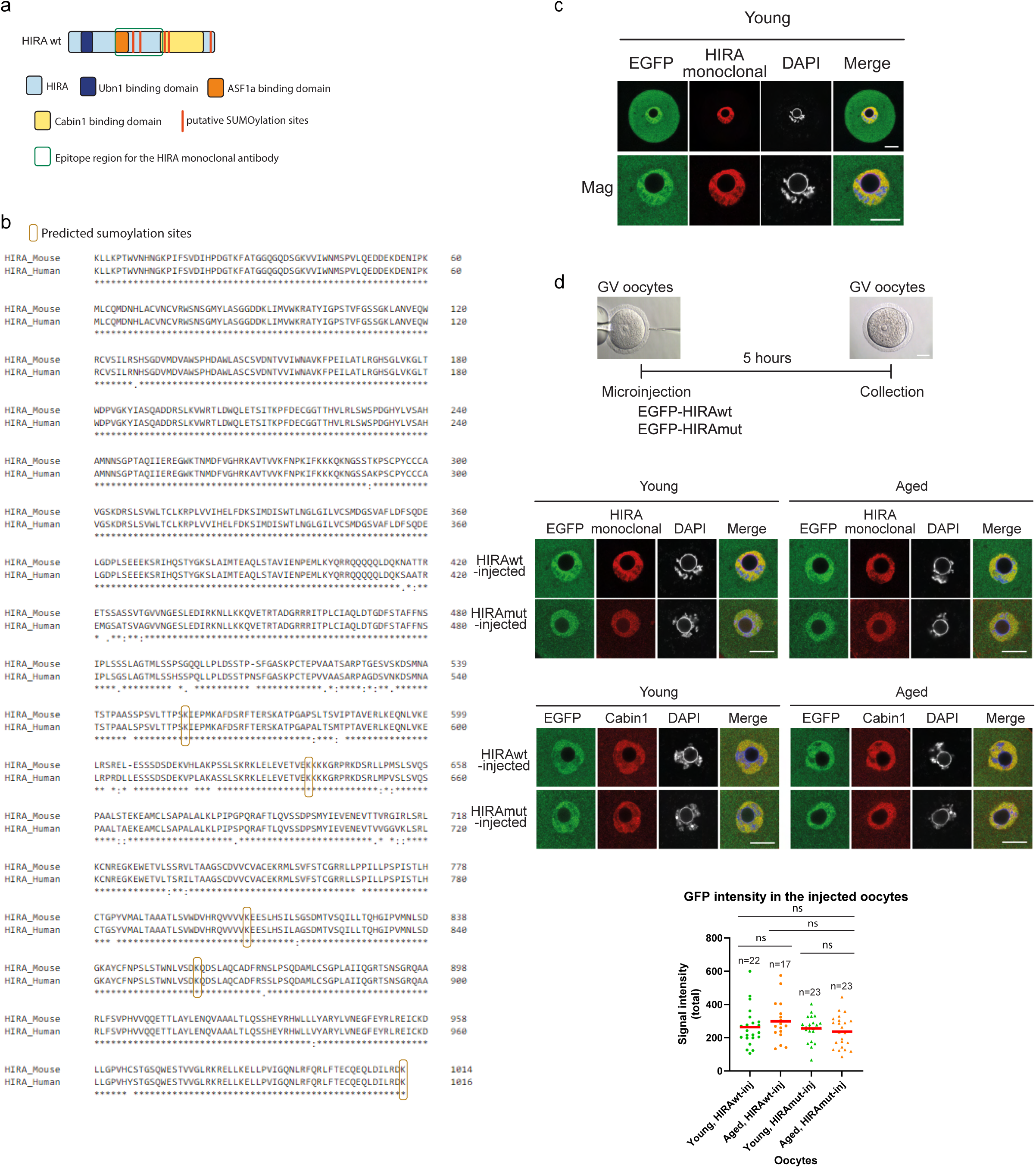
Microinjection of wt and mutated HIRA. a. Schematic diagram of HIRA protein. The red lines depict putative SUMOylation sites. b. Amino acid sequence alignment of mouse and human HIRA shows high level of conservation. The epitope region for the HIRA monoclonal antibody is 421-729aa of mouse HIRA. Predicted SUMOylation sites are shown in orange. c. Validation of the EGFP-HIRA localisation in young oocytes. IF signals show the same localisation profile as endogenous HIRA. Scale bar 20μm. d. Microinjection scheme of wt and mutated HIRA into GV oocytes. Localisation of EGFP-HIRA and Cabin1 in the EGFP-HIRAwt- and EGFP-HIRAmut-injected oocytes (total protein, no pre-extraction). Scale bar 20μm.

**Extended Data Figure 7.**
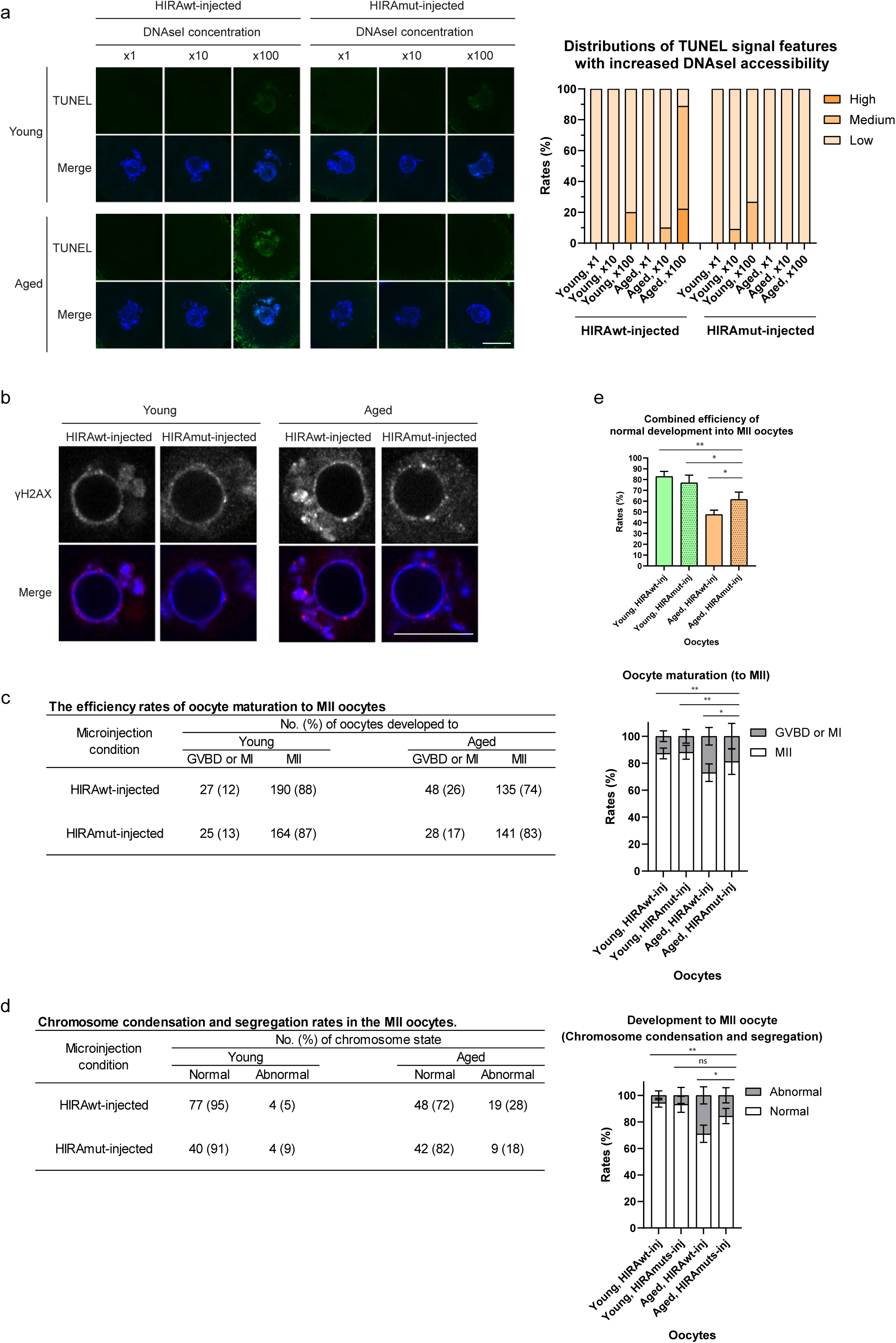
Mutation of HIRA SUMO sites restores chromatin and DNA damage, oocyte maturation, and chromosome condensation and segregation\. a. Z-Stacked images of TUNEL assay following treatment with different concentrations of DNAseI (left). The bar graph shows distribution of different signal intensities (right). Scale bar 20μm. b. High magnification of the γH2AX signal in HIRAwt- and HIRAmut- injected oocytes at SN stage, related to Figure 5d. Scale bar 20μm. c. The number (%) of oocytes developed to MII stage, related to Fig. 5e. Bar shows SD. d. The number (%) of MII oocytes with correct chromosome condensation and segregation, related to Fig. 5f. Bar shows SD. e. Combined efficiency of oocyte maturation and normal chromosome segregation. *p < 0.05. **p < 0.01.

**Extended Data Figure 8.**
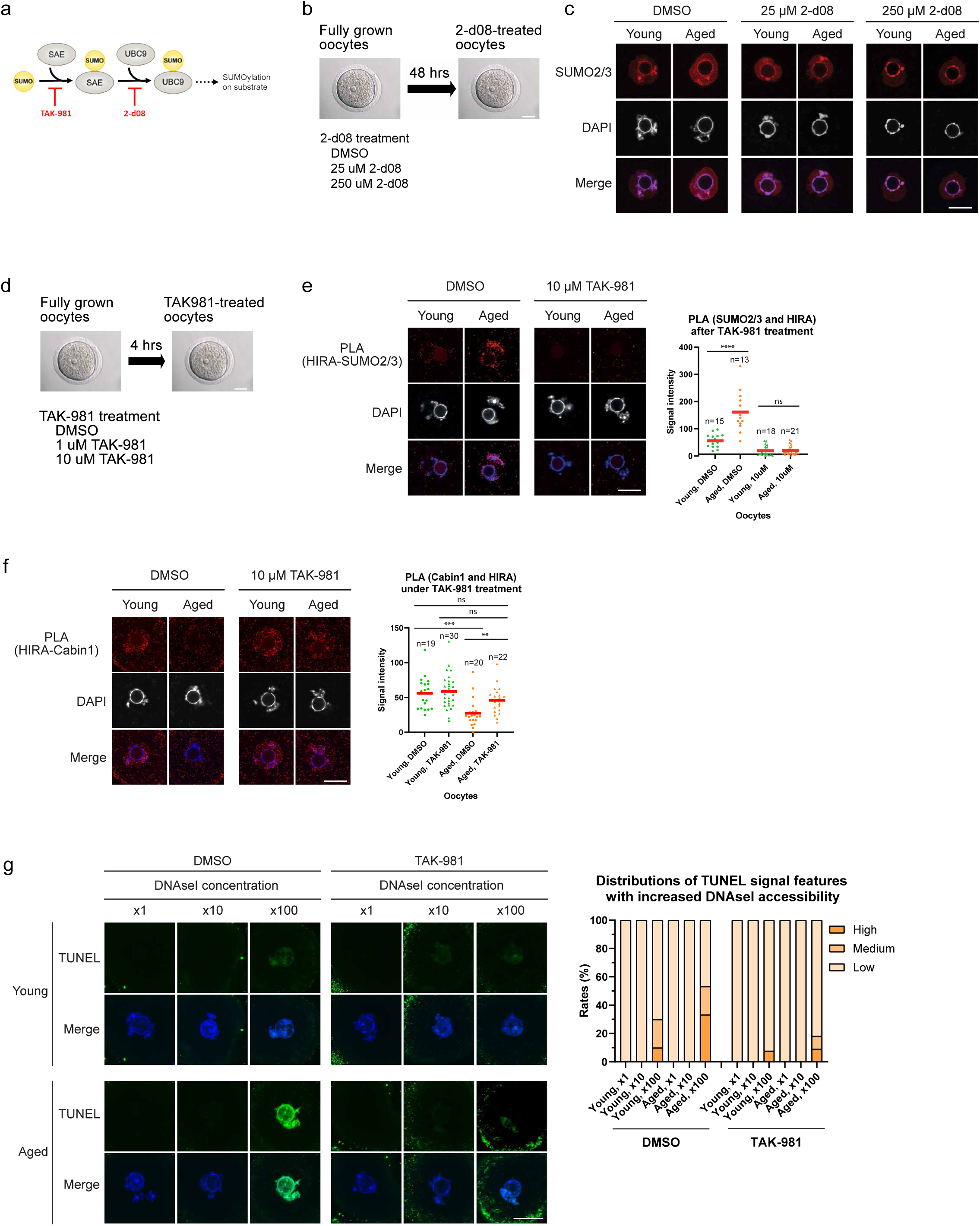
The effect of TAK-981 on oocyte maturation and chromosome condensation and segregation. a. SUMOylation pathway. TAK-981 and 2-d08 are inhibitors of SAE and UBC9, respectively. b. Scheme of 2-d08 treatment in GV oocytes. c. The effect of 48 hours’ treatment with 2-d08 on SUMO2/3 signals. Chromatin localised SUMO2/3 signal persisted in all conditions. Scale bar 20μm. d. Scheme of TAK-981 treatment of GV oocytes. e. PLA signal shows the reduction of SUMO2/3 accumulation on HIRA in the aged oocytes following TAK-981 treatment. ****p < 0.0001, ns – not significant. Scale bar 20μm. f. PLA documents that TAK-981 treatment improves HIRA-Cabin1 interaction in aged oocytes. n=number of analysed oocytes, ***p < 0.001, **p < 0.01, ns – not significant. Scale bar 20μm. g. TUNEL assay following treatment with different concentrations of DNAseI. The bar graph shows distribution of signal intensities. Scale bar 20μm.

**Extended Data Figure 9.**
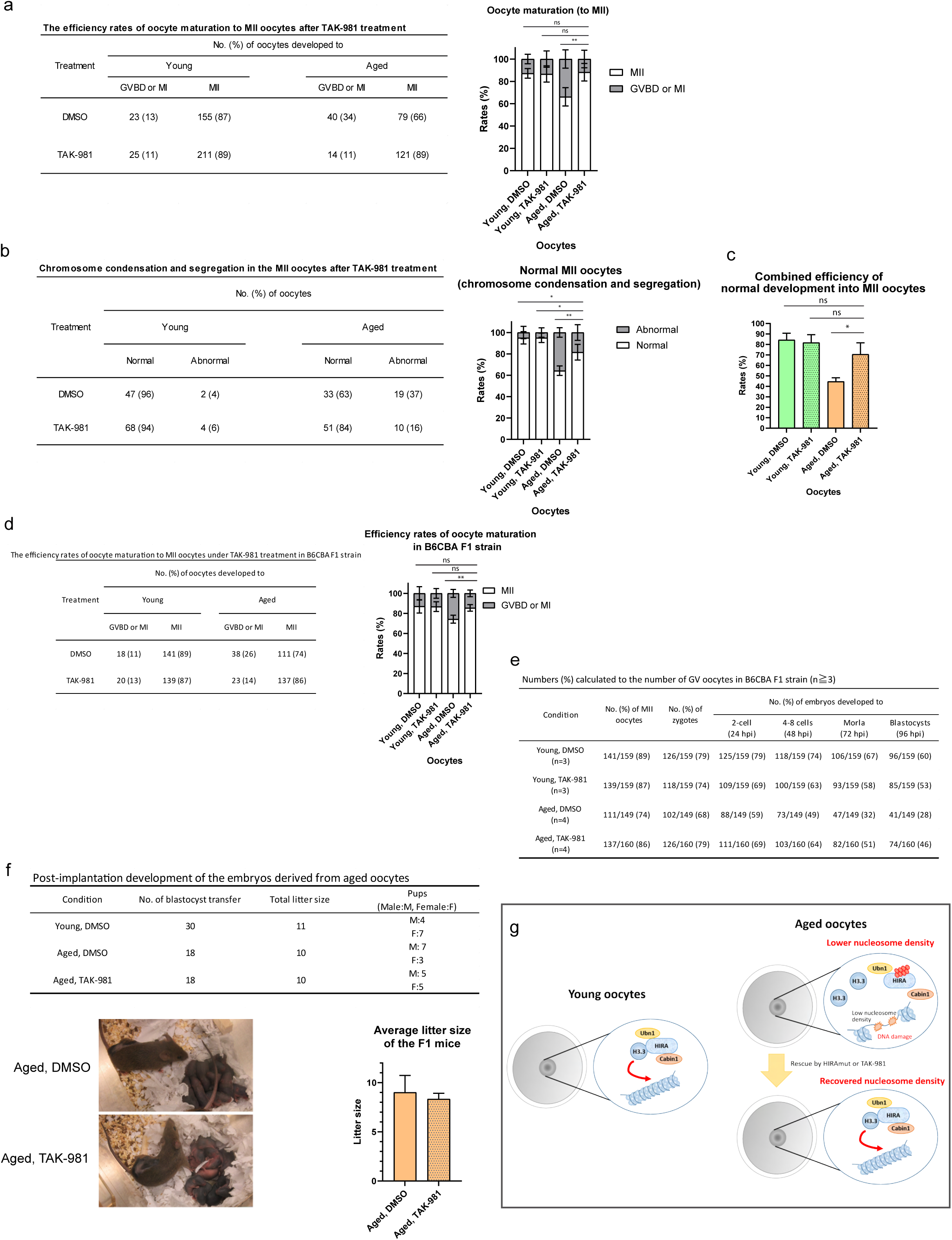
Effect of TAK-981 on chromosome organisation and developmental potential of aged mouse oocytes. a. The number (%) of oocytes developed to MII stage following TAK-981 treatment, related to Fig. 6f. **p < 0.01, ns – not significant. Bar shows SD. b. The number (%) of oocytes with correct chromosome condensation and segregation at MII stage following TAK-981 treatment, related to Fig. 6g. **p < 0.01, *p < 0.05. Bar shows SD. c. Combined efficiency of oocyte maturation and normal chromosome segregation. *p < 0.05, ns -not significant. d. The efficiency of oocyte maturation to MII stage in B6CBA F1 mice. In vitro maturation was performed using the DMSO- or TAK-981- treated fully grown oocytes. **p < 0.01, ns - not significant. Bar shows SD. e. The number (%) of oocytes/embryos developed to MII stage/blastocysts following TAK-981 treatment, related to Fig. 6h. Each rate was calculated by dividing the number of oocytes/embryos by the number of collected oocytes from ovaries. f. TAK-981 treated oocytes give rise to healthy normal pups (top). Normal litter size and normal fertility were observed. The bar graph shows normal fertility of the pups derived from aged oocytes treated with TAK-981 (bottom). Bar shows SD. Three biological replicates for mating were performed. g. Model: chromatin integrity in oocytes is dependent on the HIRA driven H3.3 incorporation. Efficient H3.3 incorporation is attenuated in aged oocytes due to SUMO modification of HIRA leading to the disruption of HIRA-Cabin binding. Microinjection of HIRA that cannot be SUMO-modified or blocking SUMO pathway using small molecule inhibitors restores H3.3 incorporation and developmental potential in aged oocytes.

